# Neural tracking of linguistic speech representations decreases with advancing age

**DOI:** 10.1101/2022.07.29.501978

**Authors:** Marlies Gillis, Jill Kries, Maaike Vandermosten, Tom Francart

**Affiliations:** Experimental Oto-Rhino-Laryngology, Department of Neurosciences, Leuven Brain Institute, KU

**Keywords:** neural tracking, speech processing, lifespan, aging, linguistic processing, speech understanding

## Abstract

**Background:** Older adults process speech differently, but it is not yet clear how aging affects different levels of processing natural, continuous speech, both in terms of bottom-up acoustic analysis and top-down generation of linguistic-based predictions. We studied natural speech processing across the adult lifespan via electroencephalography (EEG) measurements of neural tracking.

**Goals:** Our goals are to analyze the unique contribution of linguistic speech processing across the adult lifespan using natural speech, while controlling for the influence of acoustic processing. In particular, we focus on changes in spatial and temporal activation patterns in response to natural speech across the lifespan.

**Methods:** 52 normal-hearing adults between 17 and 82 years of age listened to a naturally spoken story while the EEG signal was recorded. We investigated the effect of age on acoustic and linguistic processing of speech. Because age correlated with hearing capacity and measures of cognition, we investigated whether the observed age effect is mediated by these factors. Furthermore, we investigated whether there is an effect of age on hemisphere lateralization and on spatiotemporal patterns of the neural responses.

**Results:** Our EEG results showed that linguistic speech processing declines with advancing age. Moreover, as age increased, the neural response latency to certain aspects of linguistic speech processing increased. Also acoustic neural tracking (NT) decreased with increasing age but in contrast to linguistic processing, older subjects showed shorter latencies for early acoustic responses to speech. No evidence was found for hemispheric lateralization in neither younger nor older adults during linguistic speech processing. Most of the observed aging effects on acoustic and linguistic processing were not explained by age-related decline in hearing capacity or cognition. However, our results suggest that the effect of decreasing linguistic neural tracking with advancing age at word-level is likely more due to an age-related decline in cognition than a robust effect of age.

**Conclusion:** Spatial and temporal characteristics of the neural responses to continuous speech change across the adult lifespan for both acoustic and linguistic speech processing. These changes may be traces of structural and/or functional change that occurs with advancing age.

**Highlights:**

- With increasing age, linguistic neural tracking of continuous speech decreases.
- With increasing age, the processing speed of linguistic aspects of speech slows down.
- Aging-related changes in word-level linguistic processing are affected by cognition.
- With advancing age, tracking of speech acoustics decreases in the right hemisphere.
- Older adults displayed earlier neural responses to speech acoustics.

## 1 Introduction

Aging goes hand in hand with physical and behavioral change. The brain is not spared and undergoes micro- and macrobiological alterations in structure and function. Processing speech and deducing meaning from it is a complex task that the human brain manages in a continuous manner at a fast pace. At an older age, speech comprehension seems to be largely preserved (Burke and Shafto, 2008; Shafto and Tyler, 2014), at least in optimal conditions, yet using EEG, researchers have found less efficient speech processing in older adults (Federmeier et al., 2003) and evidence for differential recruitment of neural resources (Brodbeck et al., 2018b). So far, it has not been investigated whether the observed age effects on speech processing are driven by differences in lower-level acoustic and/or higher-level linguistic speech processing. In this study, we explore the effects of age on neural responses to natural, continuous speech processing using EEG-based NT. NT is the phenomenon whereby the brain time-locks to specific aspects of the incoming speech (Brodbeck and Simon, 2020, for a review,). This method allows to explore aging effects separately for bottom-up and top-down aspects of speech processing.

### 1.1 Age-related changes in acoustic speech processing

The acoustic analysis in the primary auditory cortex consists of analyzing fundamental frequency, formant transitions and amplitude modulations in the speech envelope (Binder, 2016). By using EEG or magnetoencephalography (MEG), the neural response to auditory stimuli can be examined by averaging the signal over a large number of repetitive stimuli (i.e., event-related potentials (ERP)). Cortical auditory evoked potentials (CAEP), such as the P1-N1-P2 complex, reflect sound detection and encoding in the auditory cortex (Martin et al., 2008; Harris, 2020). In clinically normal-hearing older adults, CAEP peak amplitudes and latencies (N1 and P2) are increased compared to younger adults (McCullagh and Shinn, 2013; Tremblay et al., 2002, 2003). This has been related to decreased behavioral performance in older adults during a phoneme discrimination task (Tremblay et al., 2002, 2003). Moreover, advancing age brings along sensorineural hearing loss (presbyacusis), primarily in higher frequencies. This also affects neural responses to sounds (for a review see Cardin, 2016). In the current study, we investigate data from normal-hearing older adults.

In addition to differences in temporal patterns, evidence suggests that older adults may also use different neural resources than younger adults for the acoustic analysis. More specifically, younger adults typically display a right preference of the primary auditory cortex for processing slow modulation frequencies at 4 Hz (i.e., a modulation frequency that coincides with the rate of syllables in speech), yet older adults show a greater reliance on the left hemisphere (Farahani et al., 2020; Goossens et al., 2016).

These findings are based on artificially created experimental stimuli (e.g., repetition of isolated linguistic units) and therefore not an ideal approximation of real-life communication. NT of connected speech presents a novel way to analyze neural responses in more natural listening conditions. When using the envelope as speech representation, increased neural tracking has been found in older adults compared to younger adults in response to a narrative (Presacco et al., 2016; Brodbeck et al., 2018b; Decruy et al., 2019). The envelope consists of the slow amplitude modulations of speech. Envelope tracking mainly characterizes lower-level acoustic processing, but it has also been shown to be modulated by speech intelligibility and higher-order speech-specific processing (Peelle et al., 2013; Vanthornhout et al., 2018; Prinsloo and Lalor, 2020; Broderick et al., 2019). As of yet, it is not clear whether the observed age differences in envelope tracking are driven by processing of acoustics or of higher-order aspects. Brodbeck et al. (2018b) found that enhanced envelope tracking in older adults arises early on in areas inferior and lateral to the left primary auditory cortex. However, it is not yet clear how this result can be interpreted in terms of underlying functions. Put differently, which aspects of speech do older adults process differently than younger adults, acoustic and/or linguistic aspects?

### 1.2 Age-related changes in linguistic speech processing

To assign meaning to incoming speech, neural signals related to bottom-up acoustic cues (e.g., phoneme and word onsets) interact in a parallel fashion with top-down processes (e.g., generating predictions about linguistic units) (Federmeier, 2007; Kuperberg and Jaeger, 2016; Brodbeck et al., 2021a; Gwilliams and Davis, 2022; Hamilton et al., 2021). ERP research has found the N400 peak to be a neural correlate of semantic information, specifically its activation and its integration into the sentence context (Kutas and Federmeier, 2011; Nieuwland et al., 2020). This negative N400 response usually appears between 200 and 600 ms and is largest over centro-parietal electrodes (Kutas and Federmeier, 2011). In older adults, studies have consistently reported decreased N400 amplitudes and temporally delayed responses compared to younger adults (Kutas and Iragui, 1998; Woodward et al., 1993; Federmeier and Kutas, 2005; Tiedt et al., 2020; Gunter et al., 1992; Federmeier et al., 2002). This effect has been ascribed to a declining efficiency in older adults to generate predictions based on sentence-level context information (i.e. if a word is semantically fitting the previous words in the sentence) (Tiedt et al., 2020). However, this finding is difficult to unify with behavioral results, which generally report preserved or increased semantic and world knowledge in the elderly (Shafto and Tyler, 2014; Spreng and Turner, 2019) as well as a greater sensitivity to linguistic context (Tiedt et al., 2020). A possible explanation for these rather conflicting results may be that elderly make use of different strategies and neural structures to maintain speech comprehension at a functioning level (Tiedt et al., 2020; Reuter-Lorenz and Cappell, 2008). Indeed, evidence suggests that older adults present a more bilaterally organized activation pattern for processing linguistic aspects of speech, whereas younger adults rely more on the left hemisphere (Diaz et al., 2016; Wlotko et al., 2010; Wierenga et al., 2008; Grossman et al., 2002; Wingfield and Grossman, 2006). This finding is yet to be tested via EEG-based NT.

Although EEG research using naturally spoken stories has traditionally focused on bottom-up processing aspects of speech, recent NT studies demonstrate its potential to gain insights into higher-order language processing by analyzing the neural response related to the amount of linguistic information conveyed by a phoneme or word (Brodbeck et al., 2018a; Alday, 2019; Brennan and Hale, 2019; Weissbart et al., 2019; Broderick et al., 2021; Mesik et al., 2021; Shain et al., 2020; Gillis et al., 2021; Heilbron et al., 2022; Di Liberto et al., 2019). The neural response to the surprisal of a word or phoneme represents the mismatch between top-down predicted and bottom-up perceived linguistic units. Hence, the more unexpected or surprising the linguistic unit is, the larger the prediction error is (Weissbart et al., 2019; Friston, 2010). The relationship between such a surprisal model and the EEG signal can be studied via NT. This way, Broderick et al. (2021) found longer latencies of the neural response to word surprisal in older adults. However, the study by Mesik et al. (2021) did not confirm these results of Broderick et al. (2021). In both studies, the negative peak appeared during the same time window as the N400 ERP and displayed a similar centro-parietal topography. From functional magneto-resonance imaging (fMRI) research measuring the hemodynamic activity in response to a story, the word surprisal peak seems to be specific to the language network (Shain et al., 2020). These EEG studies investigating linguistic speech processing in older adults did not control the influence of acoustic processing in their model and therefore, the results may be biased by age-related effects in bottom-up acoustic processing (Gillis et al., 2021).

### 1.3 Current study

In this study, we investigated age-related spatiotemporal neural changes in acoustic and linguistic aspects of speech processing respectively. Furthermore, we analyzed the unique contribution of linguistic speech processing in elderly by controlling for acoustic and lexical segmentation representations (Gillis et al., 2021). To do so, we investigated EEG responses to natural, connected speech in neurologically healthy, normal-hearing adults ranging from 17 to 82 years of age. We investigated neural tracking of speech using a forward modeling approach. This method describes how and how well the brain responds to speech representations in a time-locked fashion, respectively by the temporal response function (TRF) (”how”) and prediction accuracy (”how well”). The prediction accuracy is a measure of neural tracking (NT), i.e., the better the brain tracks the incoming speech, the higher the prediction accuracy will be. In order to investigate the unique contribution of linguistic speech processing, we analysed the increase in prediction accuracy when linguistic representations are added to a model consisting of acoustic and lexical segmentation representations. This increase in prediction accuracy, which can be attributed to the linguistic representation, is further denoted as linguistic tracking, i.e., the tracking of linguistic aspects of the incoming speech when controlling for acoustics and lexical segmentation. We here solely focused on linguistic aspects related to generating predictions about upcoming speech, namely phoneme surprisal, phoneme entropy, word surprisal and word frequency.

Our first research question concerned the effects of age on the magnitude of NT separately for acoustic and linguistic features. Given that age, hearing and cognition are interrelated, we performed additional analyses to examine whether any observed age effects on the acoustic and linguistic aspects of speech are mediated by decline in hearing or cognitive performance respectively. Additionally, spatial pattern differences of the magnitude of NT across age were investigated in predefined regions of interest and in the hemispheres. The second research question regarded temporal pattern changes across age, which were analyzed by means of TRF peak latencies. Again, we investigated whether the effect of age on the latency was mediated by hearing capacity or cognition. Additionally, the topographies of the TRF peaks were investigated.

For acoustic speech processing, we hypothesized to find increased NT with increasing age (Presacco et al., 2016; Brodbeck et al., 2018b; Decruy et al., 2019). This effect was expected to be strongest in the auditory cortex, situated in temporal regions of the brain (Cardin, 2016). Moreover, based on previous findings, we hypothesized that there would be stronger left-hemispheric lateralization in older than in younger adults for the acoustic model (Brodbeck et al., 2018b). Regarding temporal patterns, we hypothesized longer latencies with advancing age, given that most studies investigating CAEP reported longer latencies in older adults. Additionally, we hypothesized that this age effect is driven by hearing capacity: with increasing age, hearing capacity decreases, which might affect neural speech processing. Previous literature showed that adults with hearing loss showed higher acoustic NT and later neural response latencies compared to normal hearing adults (Gillis et al., 2022), which supports our hypothesis.

For linguistic speech processing, we hypothesized a decrease in linguistic NT with advancing age. Although Broderick et al. (2021) and Mesik et al. (2021) reported diverging results comparing linguistic NT between younger and older adults, ERP studies consistently suggested that older adults show smaller and later N400 responses. Although the link between NT and ERP amplitudes is not well established, we assume that language processing changes across the lifespan, which would affect both the NT as well as the spatial and temporal characteristics of the neural responses. Based on ERP (Kutas and Federmeier, 2011) and NT (Broderick et al., 2021; Mesik et al., 2021) literature, we expected the age-effect to be strongest over the centro-parietal region. Concerning hemispheric lateralization, we hypothesized more symmetrical NT in older than in younger adults for the linguistic speech processing (Diaz et al., 2016; Wlotko et al., 2010). Regarding temporal dynamics, the TRF latencies of linguistic speech representations were expected to be longer in older adults, based on the results from Broderick et al. (2021), who found increased latencies in older adults in response to word surprisal. Moreover, we hypothesized that the effect of age is mediated by cognitive factors: with increasing age, the brain undergoes age-related changes which goes hand in hand with a decline in cognition. We investigated these age-related changes by analysing the changes in the spatial and temporal dynamics of the neural responses. We hypothesized that cognition would impact linguistic NT and the neural response latencies to linguistic representations.

## 2 Material and Methods

We investigated acoustic and linguistic speech processing across the adult life span. For each type of speech processing, we investigated two outcome variables: *the prediction accuracy*, also denoted as neural tracking (NT), to investigate age-trends and lateralization across age, and the *TRF pattern* to study how the spatial and temporal characteristics of the neural responses to the speech representations change with age. We used a subset of the data collected in the study by Decruy et al. (2019). In comparison to Decruy et al. (2019), we investigated NT of a more extensive set of speech representations covering acoustic and linguistic aspects of the speech, rather than just the speech envelope investigated in Decruy et al. (2019).

### 2.1 Participants

We analyzed the data of 52 normal-hearing adults. Age varied between 17 and 82 years old. All participants had a normal hearing, confirmed with pure tone audiometry (no hearing threshold exceeded 30 dB HL at all octave frequencies from 125 to 4000 Hz; defined identical to Decruy et al. (2019)). The average of these measured hearing thresholds is denoted as pure tone average (PTA) (octave frequencies from 125 to 4000 Hz; mean*±* std = 12.1*±* 5.08; the distribution of PTA across age is visualized in fig. 2; the raw thresholds are shown in the supplementary material in figure S.1). Same as in Decruy et al. (2019), we excluded individuals with a low cognitive score, i.e., Montreal Cognitive Assessment (Nasreddine, 2004) *<* 26/30, and individuals with a learning disability or medical history of head concussions. The data of two participants of the original dataset, collected by Decruy et al. (2019), were discarded: one participant was discarded because the stimulus was presented to the left instead of the right ear; another participant was missing an EEG electrode. The experiments were carried out following a protocol accepted by the KU Leuven Ethical Committee. All participants signed an informed consent form. For more details regarding the cognitive screenings, we refer to Decruy et al. (2019).

### 2.2 Cognitive testing

Decruy et al. (2019) conducted two cognitive tests. Cognitive performance was assessed using the Stroop color-word test (SCWT) (Hammes, 1978), measuring inhibition, and the reading span test (RST), measuring working memory (van den Noort et al., 2008; Vercammen et al., 2017).

During the RST, the participants were instructed to read sets of 20 sentences aloud. After each set, the participants recalled as many as possible final words of sentences of the set. After three sets of sentences, the working capacity is represented by the amount of correctly recalled final words of 60 sentences as possible (resulting in a score of 60). A higher RST outcome represents better working memory performance.

For the SCWT, participants were shown three cards containing respectively color names, colored rectangles and color names printed in incongruent colors. For each card, the task instruction was to name or read the color shown on the card as fast as possible. For the card containing the color names printed in incongruent colors, the task consisted of naming the color of the word and inhibiting to read the color name. The SCWT score is calculated as the difference in response time (in milliseconds) between the third and the second testing condition. Therefore, a higher SCWT score represents a higher difference in response time and thus a decreased inhibition performance.

For a more extensive explanation of how the cognitive tests were carried out, we refer to Decruy et al. (2019).

### 2.3 EEG testing

During the EEG experiment, the participants listened to a story in Flemish, titled Milan and written by Stijn Vranken (male speaker). The story was presented in silence. It lasted 12 minutes and was narrated by the author. The neural responses associated with listening to this story were investigated to analyse acoustic and linguistic NT.

The stimuli were presented to the right ear at an intensity of 55 dB SPL (A-weighted). During the stimulus presentation, EEG data was collected using a BioSemi Active Two system (Amsterdam, Netherlands) with 64 EEG electrodes. For more details on the experimental protocol, we refer to Decruy et al. (2019).

### 2.4 EEG processing

Firstly, to decrease the processing time, the EEG recording with a sampling frequency of 8192 Hz was downsampled to 256 Hz. Subsequently, we filtered the EEG using a multi-channel Wiener filter (Somers et al., 2018) to remove artifacts due to eye blinks and referenced the EEG to the common-average. We filtered the data between 0.5 and 25 Hz using a Chebyshev filter (Type II with an attenuation of 80 dB at 10% outside the passband). Lastly, an additional downsampling step was done to reduce the sampling rate to 128 Hz. Preprocessing of the EEG was performed in MATLAB using custom scripts (MATLAB, 2016).

### 2.5 Predictors

The speech representations used in this study are based on the findings of Gillis et al. (2021). We divided the speech representations into two categories (1) acoustic representations, and (2) linguistic speech representations. The acoustic representations resembled the continuous acoustic energy of the presented speech material, while the linguistic speech representations denote to what extent a phoneme or word can be predicted based on the context. The linguistic representations at the word level were calculated for each word, independent of word class. Similar to Brodbeck et al. (2018a), the linguistic representations at the phoneme level were modelled excluding the first phoneme.

The acoustic speech representations were calculated from the speech stimulus. First, to account for the filtering of the ER-3A insert earphones, the speech stimulus was low-pass filtered below 4000 Hz (zero-phase low-pass FIR filter with a hamming window of 159 samples). The ***spectrogram*** representation was calculated using the Gammatone Filterbank Toolkit 1.0 (Heeris, 2014) with center frequencies between 70 and 4000 Hz with 256 filter electrodes and an integration window of 0.01 second. This toolkit was used to calculate a spectrogram representation based on a series of gammatone filters inspired by the human auditory system’s structure (Slaney, 1998). The resulting 256 filter outputs were averaged into eight frequency bands (each containing 32 gammatone filter outputs). Additionally, each frequency band was downsampled to the same sampling frequency as the processed EEG, namely 128 Hz. Subsequently, the ***acoustic onsets*** representation was calculated as the half-wave rectification of the spectrogram’s derivative.

Linguistic speech representations were obtained at the phoneme and word level. ***Phoneme surprisal*** and ***phoneme entropy*** were used as two linguistic speech representations at the phoneme level (Brodbeck et al., 2018a). Both representations were derived from the activated cohort of words, given a certain sound (e.g., when hearing the sound *sp*, the activated cohort of words include words like *speed, spam, special*, etc.). Phoneme surprisal represents the surprisal of a phoneme. It was calculated as the negative logarithm of the phoneme probability in the activated cohort. In contrast, phoneme entropy reflects the degree of competition between the words congruent with the phonemic input at a given point during a word. It was calculated as the Shannon entropy of the words in the active cohort. A higher value indicated that more phonemes have a high probability of occurring as a subsequent phoneme. These representations at the phoneme level were calculated using the SUBTLEX-NL database (Keuleers et al., 2010) and a custom pronunciation dictionary maintained in our lab. Gillis et al. (2021) showed that the EEG response to these representations is characterized by a topography with central negativity around 250 ms.

Concerning linguistic representations at the word level, ***Word surprisal*** was calculated, which reflects the surprisal of a word given the four previous words. It was calculated as the negative logarithm of the conditional probability of the word, given the four preceding words. ***Word frequency*** is calculated as the negative logarithm of the unigram probability. Note that due to this negative logarithm, frequent words are projected onto lower values. These representations at the word level were calculated using the 5-gram model by Verwimp et al. (2019) using the corpora *Corpus of Spoken Dutch* (Oostdijk et al., 2000) and a database of subtitles. The response to word surprisal is characterized by the topography with central negativity around 400 ms (Weissbart et al., 2019; Gillis et al., 2021). This response resembles the N400-effect in ERP responses.

To control for the early acoustic response, we included speech representations for phoneme and word onsets, i.e., lexical segmentation. The phoneme and word onset timings were determined by using a forced aligner (Duchateau et al., 2009). The phoneme and word onset representations describe the onsets with a hot-one encoding: 1 at the time of onset while all other values are 0. As the response to phoneme and word onsets is not purely acoustic, we did not include these representations when evaluating acoustic NT. When investigating linguistic tracking of speech, controlling for acoustic and lexical segmentation representations is necessary (Gillis et al., 2021). If this step is not considered, a linguistic representations might be significantly tracked due to correlations with acoustic or lexical segmentation representations.

### 2.6 Neural tracking

The forward modelling approach provides insight into the brain’s response to speech in two ways: a measure of NT, the prediction accuracy, and a temporal response function (TRF), which characterizes the brain’s temporal response to the speech representations. The TRF models how the brain responds to these speech representations. Subsequently, the TRF is used to predict the EEG by convolving it with the speech representations. Correlating the predicted responses with the measured EEG results in a prediction accuracy for each EEG electrode. These prediction accuracies are a measure of NT: the higher the prediction accuracy, the more the brain tracks the speech representations.

We used the Eelbrain toolbox to estimate the TRF and to calculate the prediction accuracies (Brodbeck et al., 2021b). We estimated the TRF using boosting (David et al., 2007), in a cross-validation approach with ten partitions and an integration window between 0 and 600 ms (using selective stopping based on the *£*_2_-norm and Hamming window of 50 ms as basis function). Additionally, we calculated the prediction accuracy in three different testing windows that each characterize a different type of response:

- 30 to 160 ms: the early acoustic response like N1 in the spectrogram
- 160 to 300 ms: a later acoustic response like the P2 in the spectrogram and phoneme recognition which occurs around 250 ms as observed in the TRF to phoneme surprisal and phoneme entropy
- 300 to 600 ms: linguistic processes like the N400 response to word surprisal

**Figure 1:**
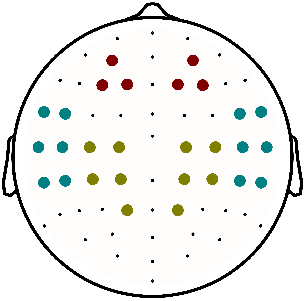
Different electrodes selections: frontal (red), centro-parietal (green) and temporal (blue).

These testing windows were determined based on the average TRF across all participants. Outside the testing windows, we set the TRF to zero. This adapted TRF was convolved with the speech representations to obtain the prediction accuracy based on the lags specified in the testing window. We opted to set the TRF, trained on the integration window from 0 to 600 ms, partially to zero over retraining the TRF in different integration windows. This was done to prevent edge effects, i.e., high values at the beginning and end of the window, which occur due to autocorrelations in the speech.

We classified NT into two different types: (1) acoustic NT and (2) linguistic NT. Acoustic NT denoted the prediction accuracies of a model using acoustic speech representations of the speech (i.e., spectrogram and acoustic onsets). Linguistic NT was calculated as the prediction accuracy increase when linguistic speech representations are included in the model on top of acoustic speech representations and lexical segmentation representations as phoneme and word onsets. In more detail, we subtracted prediction accuracies from the model that included acoustic and lexical segmentation representations from those obtained with the model that included acoustic, lexical segmentation, and linguistic speech representations. The difference between these models thus constitutes the added value of linguistic representations over and beyond acoustic and lexical segmentation properties. Therefore, if the prediction accuracy is significantly higher than zero, the linguistic speech representations explain unique information over and beyond the response to the acoustic signal or the onset of a word or phoneme. If acoustic and lexical segmentation representation are not taken into account, spurious tracking might be observed leading to biased results.

### 2.7 Regions of interest

We investigated the effect of age on acoustic NT and linguistic NT averaged across all EEG electrodes. However, this approach can average out sublte and local effects. Therefore, we also investigated acoustic NT and linguistic NT in three a priori-chosen electrode selections, i.e., a frontal electrode selection, a centro-parietal electrode selection, and a temporal electrode selection (visualized in fig. 1). The frontal electrode selection was based on previous research showing that acoustic neural tracking displays the highest magnitude over these sensors (Lesenfants et al., 2019). The temporal electrode selection was on the one hand hypothesis-driven, given that the temporal cortex is largely involved in auditory processing, and on the other hand based on previous research, i.e., Brodbeck et al. (2018b). The centro-parietal electrode selection was based on results from ERP studies showing that the N400 component is most prominent over centro-parietal sensors and on results from Gillis et al. (2021), showing that linguistic features have a prominent neural response over centro-parietal sensors.

### 2.8 Determination of lateralization

To determine whether there was an age-dependent hemisphere effect, we averaged the prediction accuracy across respectively right or left-side electrodes, excluding central electrodes. This analysis was performed across all EEG electrodes and for the individual ROIs.

### 2.9 Determination of peak latency

To determine the peak latency, we defined a time window for each peak of interest, based on the average TRF across all participants, wherein we investigated the amplitude and latency. We broadened these time windows on each end with 30 ms to generalize these regions across all participants because the average TRF is biased towards 50-year-olds as age was not equally distributed across all participants (see age-distribution in fig. 2). The resulting time windows are summarized in table 1. Within each time window, we determined the root mean square (RMS) of the TRF across electrodes for each time lag. The RMS is higher when many electrodes have high activity, indicating the possibility of a peak. In each local maximum of the RMS, we correlated the peak topography with a template peak topography, i.e., a peak’s topography determined across all participants and averaged across the whole time window (table 1). We selected the latency of the local maximum where the peak topography had the highest positive correlation with the template peak topography. Additionally, we set a criterion that more than 30 electrodes must have the same sign as the template peak topography. If not, we discarded the peak latency and topography from the analysis. The number of peaks found per speech representation and time window is shown in table 2. To investigate whether this peak was significant, we repeated this template matching peak finding on the shuffled TRF within this time region. The significance of the peak was determined based on the 95th-percentile value of correlation between the template and the peak in the shuffled TRF of 1000 permutations. This method has been validated and compared to the manual determination of the peak latencies.

**Table 1:**
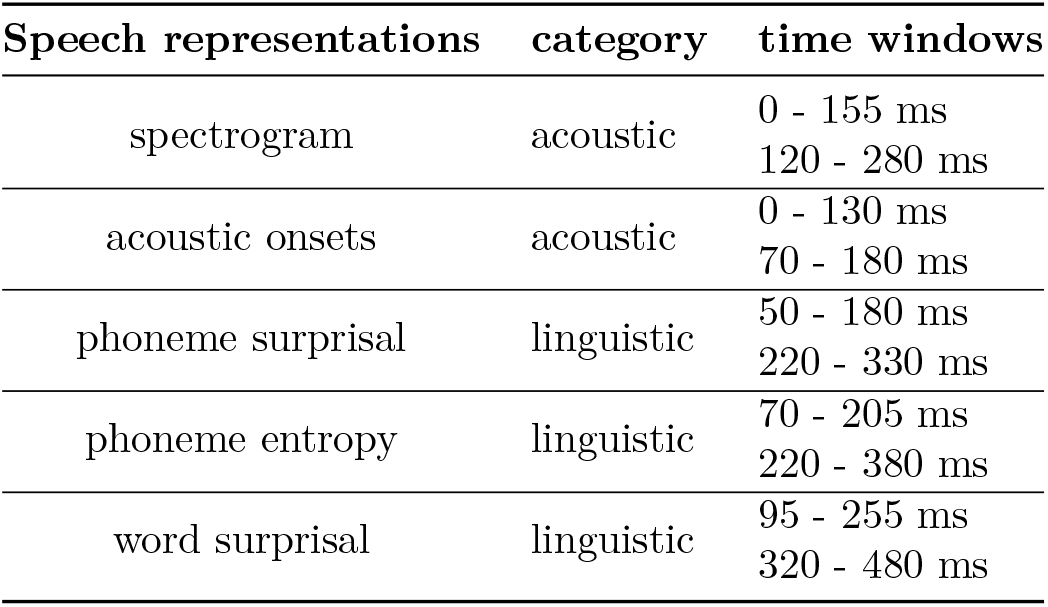
Selected time windows for each speech representation to determine the peak latencies.

**Table 2:** Percentage of peaks found per speech representation and time window.

| Speech representation | age-label | early window | late window |
| --- | --- | --- | --- |
| spectrogram | younger | 82% | 100% |
|  | middle | 68% | 100% |
|  | older | 100% | 100% |
| acoustic onsets | younger | 100% | 82% |
|  | middle | 100% | 86% |
|  | older | 93% | 100% |
| phoneme surprisal | younger | 73% | 100% |
|  | middle | 79% | 72% |
|  | older | 100% | 77% |
| phoneme entropy | younger | 69% | 82% |
|  | middle | 78% | 86% |
|  | older | 77% | 69% |
| word surprisal | younger | 55% | 100% |
|  | middle | 68% | 93% |
|  | older | 100% | 54% |
| word frequency | younger | 73% | 82% |
|  | middle | 79% | 93% |
|  | older | 92% | 54% |

In total 52 participants: 11 younger adults, 28 middle aged adults, 13 older adults

Code for this method to determine peak latencies is available online (https://github.com/exporl/TemplateMatching).

### 2.10 Statistics

The statistical analyses were carried out with Rstudio (RStudio Team, 2020), which uses R software (R Core Team, 2020). All linear models to determine age effects on the prediction accuracy, averaged across all electrodes, or peak latency were created using the lme R package (Bates et al., 2015). The model characteristics are reported with the adjusted 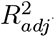-value, F-value, degrees of freedom and p-value. The estimates of the considered effect are reported with estimates, represented by *β*, standard errors, t-values and corresponding p-values.

Similarly as the linear models, the linear mixed effect models were created using the lme R package (Bates et al., 2015). These models were used to investigate the effect of age on the prediction accuracy in specific ROI and/or hemipheric lateralization. To investigate the effect of age on the prediction accuracy in the different ROI, linear mixed effects model were created with ROI as a 3-level factor variable (fixed effect) and participant as random effect (*prediction accuracy∼age+ROI+age*ROI+(1|participant)*). To investigate the effect of age on lateralization, we determined whether the effect depended on the considered hemisphere by using hemisphere as a 2-level factor, age as fixed effect, and participant as a random effect (*prediction accuracy∼age+hemisphere+age*hemisphere+(1|participant)*). In case the hemisphere and age interaction effect was significant across all electrodes, the same analysis was repeated separately for the 3 region of interest (ROI)s (*prediction accuracy in ROI∼age+hemisphere+age*hemisphere+(1|participant)*). false discovery rate (FDR) correction was applied to account for multiple comparisons (Benjamini and Hochberg, 1995).

These linear mixed effect models were fitted using using the restricted maximum likelihood method. Posthoc pair-wise comparisons were performed using the R package emmeans (Lenth et al., 2022), the Tukey contrasts were applied and corrected for multiple comparisons applying the FDR method (Benjamini and Hochberg, 1995). The model characteristics are reported with the adjusted *R*^2^*conditional*-value. Similar to the *R*^2^ for linear models, *R*^2^*conditional* represents the estimator of the explained variance by the linear mixed effect model (Barton, 2022). The estimates of the considered effect are reported with estimates, represented by *β*, standard errors, degrees of freedom, t-values and corresponding p-values.

To examine whether the direct effect of age persists after considering the correlated behavioral measures, i.e., PTA, SCWT, and RST, we applied structural equation modelling (SEM). We applied SEM using the Lavaan R package (Rosseel, 2012). This package allows estimating the indirect effect of a certain predictor through other predictors. To assess the goodness of fit, we used the criteria according to Hu and Bentler (1999): Comparative Fit Index (CFI) *>* 0.95, Tucker-Lewis Index (TLI) *>* 0.95, root mean squared error of approximation (RMSEA) *<* 0.06. For this analysis, all variables were scaled by subtracting the mean and dividing by their standard deviations. The outcomes of SEM are reported as estimates of the effect, represented by *β*, the standard error (SE), z-value, and p-value.

To investigate the effect of age on the topographies, we created two groups consisting of the older (*>* 60 years; n = 13) and younger (*<* 40 years; n = 11) participants. This division into two groups allows us to compare how the TRF patterns differed between younger and older participants. For investigating differences in topographies, we used two different methods. Firstly, an independent cluster-based permutation test proposed by Maris and Oostenveld (2007), using the Eelbrain implementation (Brodbeck, 2020). This method allows pinpointing the electrodes which drive the significant difference between both groups. However, a disadvantage of using such a test is a preset threshold for defining a cluster. Depending on the value of this threshold, the test is more specific to either weaker and widespread effects or strong and localized effects. To overcome this, we applied the method proposed by McCarthy and Wood (1985). This method compares normalized topographies, therefore discarding amplitudes effects. More specifically, the method is based on an ANOVA test for an interaction between the normalized topography value of electrode and condition (here: younger and older adults), i.e. testing whether the age label modulates the normalized response pattern. However, this method does not allow to pinpoint which electrodes drive the significant difference.

By investigating the spatial patterns of the prediction accuracies or TRF peak topographies, one can obtain useful information about the underlying neural source configuration. A change in spatial pattern implies that the underlying neural source configuration changed. Either the same neural sources are active but their relative magnitude of activation changed or different neural sources are active. Note that, the other way around, i.e. if there is no difference in spatial pattern, that this does not necessarily mean that the underlying source configuration is constant.

To investigate the effect of age on the frontal TRF pattern, we applied the cluster-based permutation test to identify whether the TRF pattern was significantly below 0.

A significance level of *α* = 0.05 was used. For the cluster-based permutation test, this significance level was divided by the amount of comparisons to correct for multiple comparisons.

## 3 Results

We evaluated the effect of age on acoustic and linguistic NT. First, we focused on the effect of age on NT by analysing the prediction accuracies (points 1.a. and 1.b.). Second, we investigated the temporal and spatial patterns of the neural response by exploring the TRF pattern (points 2.a. and 2.b.).

1.a. We identified whether age significantly affects the NT averaged across all electrodes. Additionally, we investigated whether the observed age effect on NT is mediated by other measures of age-related decline like hearing capacity and cognition.
1.b. We investigated the spatial pattern in NT within the a-priori selected ROIs and investigated hemispheric lateralization.
2.a. We characterized how age affects the temporal pattern of the neural response by analyzing the effect of age on the neural response latencies. Similarly, we analyzed whether the effect of age on the response latency is mediated through hearing capacity or cognition.
2.b. Lastly, we studied the associated topographies to investigate whether age affects the underlying neural source configuration responding to the different speech representations.

### 3.1 Effect of age on hearing capacity and cognitive measures

We evaluated the correlation between age and hearing capacity and cognitive measures (fig. 2). We observe a positive correlation between age and PTA: with increasing age, a higher PTA value is observed (Pearson’s r = 0.478, p *<* 0.001). Moreover, when evaluating the cognitive performance across the age span, we observed that age is positively correlated with the SCWT scores (Pearson’s r = 0.494, p *<* 0.001) and negatively correlated with the RST outcome (Pearson’s r = -0.445, p *<* 0.001). Both measures reflect the cognitive performance but in the opposite direction, namely, a worse cognitive performance is associated with a higher SCWT score and a lower RST outcome. Altogether the pattern of the latter two measures across age indicates that as age increases, cognitive performance declines.

**Figure 2:**
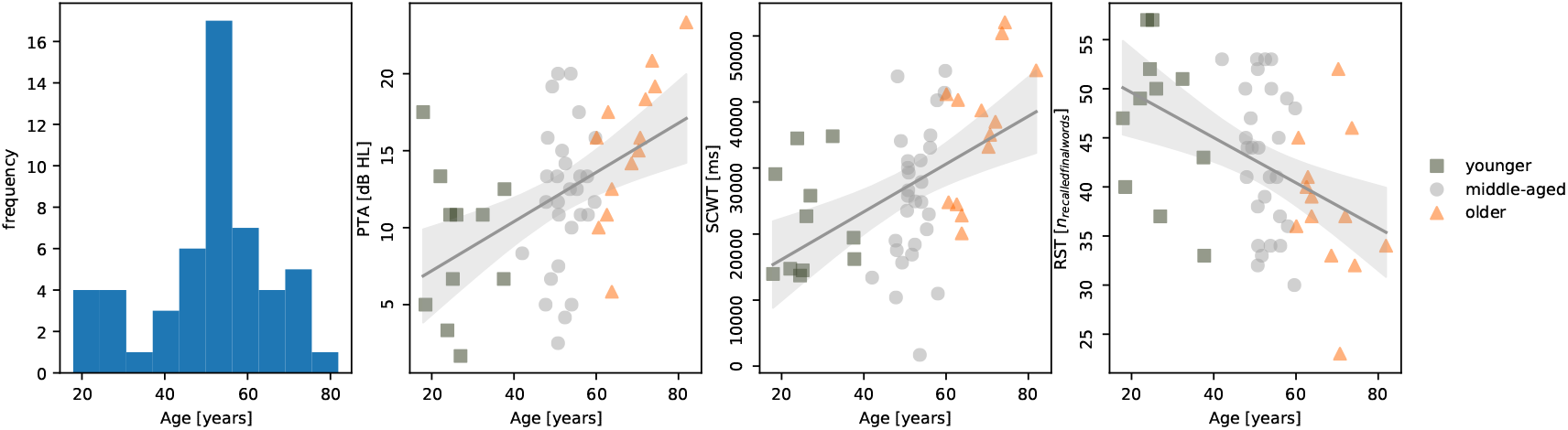
The distribution of age across the participants, pure tone average (PTA), Stroop color-word test (SCWT) scores and reading span test (RST) across age. The data points of older adults (*>* 60 years old) are represented by orange triangles, while those of the youngest adults (*<* 40 years old) are represented by green squares. All other participants, referred to as middle-aged participants, are represented by grey dots.

### 3.2 Effect of age on acoustic speech processing

#### 3.2.1 Prediction accuracy

To investigate the effect of age on acoustic NT, we investigated how the prediction accuracy, averaged across time windows (30-160 ms and 160-300 ms) and over EEG electrodes, is affected by age. Acoustic NT was obtained using the forward modelling approach with the two acoustic representations: spectrogram and acoustic onsets. Using a linear model, we did not observe a significant effect of age on the prediction accuracy of the acoustic model averaged across all electrodes (p = 0.128). Statistics are reported in table 3.

Given that local and subtle effects cannot be detected when the prediction accuracy is averaged across all EEG electrodes, we investigated whether this age effect is observed for a priori defined sets of electrodes via a linear mixed effects model. This demonstrated that acoustic NT decreases with increasing age when NT is evaluated in the 3 ROIs, i.e., main effect of age (p = 0.033). The model showed no main effect of ROIs nor an interaction effect (fig. 3.B). Notably, these results differ from the previous analysis including all 64 EEG channels. This effect can be explained as only 28 of the 64 channels were included in this analysis.

Using a SEM analysis, we verified whether this age effect persists after considering the hearing capacity, i.e., after including the PTA values, and cognitive measures (more details in section Statistics under Methods). This analysis was preformed averaged across the 3 ROIs and across the two earlier time windows. Indeed, a direct effect of age on the neural tracking of acoustic representations is observed (*β* = -0.502, p = 0.003; additional details regarding this analysis are included in the supplementary material). Moreover, this analysis indicated that the effect of age is opposite to the effect of PTA. Neural tracking decreases with increasing age, while neural tracking increases with an increasing degree of hearing loss.

**Figure 3:**
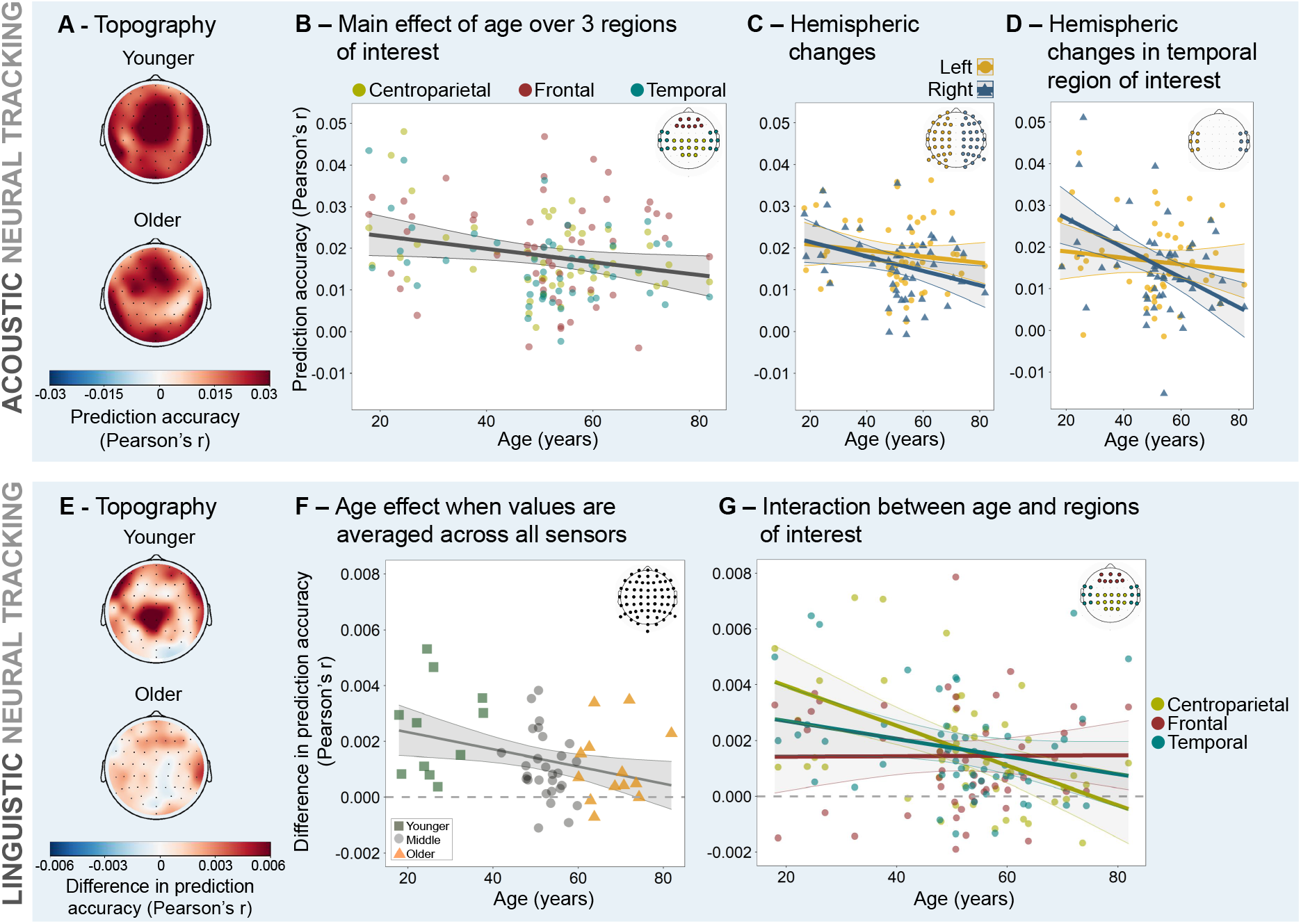
Top panel A-D: Age affects acoustic neural tracking in regions of interest and interacts with hemisphere. **A.** Average topography of younger participants below 40 years of age and older participants above 60 years of age for visualization purpose. **B.** A linear mixed effects model revealed a main effect of age over 3 ROIs, which did not differ from each other. **C.** Interaction effect between age and hemisphere when values are averaged across all left and all right hemisphere electrodes respectively. **D.** Interaction effect between age and hemisphere when values are averaged across left temporal and right temporal electrodes respectively. **Bottom panel E-G: Age affects phoneme- and word-level linguistic neural tracking. E.** Average topography of the younger participants and older participants for visualization purpose. **F.** Linear age effect when values are averaged across all electrodes. **G.** A linear mixed effects model revealed an interaction between age and ROI. Post-hoc comparisons showed that the centro-parietal ROI is significantly different from the frontal one. The insets in B, C, D, F and G show the EEG electrodes included in the analysis. All statistics are reported in table 3.

Opposite to previous literature, we did not find that with increasing age, acoustic NT increases. As we are using the same data as Decruy et al. (2019), we investigated the differences in more detail. Although Decruy et al. (2019) used a different modelling approach, we aimed to replicated the finding using the forward modelling approach with similar speech representations and frequency band as used in Decruy et al. (2019). Namely, we used a frequency band of 0.5 to 8 Hz (same filter details as mentioned in the method section) and an envelope representation (calculated as the spectrogram averaged across the eight frequency bands) to allow a better comparison to the results of Decruy et al. (2019). However, similarly as above, we did not observe any effect of age on the average prediction accuracy over all electrodes using acoustic representations (model characteristics: 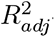 = 0.006; F = 1.32; df = 50; p = 0.256; age-effect: *β* = -1.00e-04, SE = -1.75e-04, t = -1.149, p = 0.256). Moreover, when including a quadratic age term, this quadratic term does not reach significance or improve the model fit. It is noteworthy that there is a substantial methodological difference between Decruy et al. (2019) and our results although using the same dataset. Firstly, the training paradigm is significantly different. Decruy et al. (2019) trained the model on the 12-minute story and applied it to a different speech stimulus (not investigated in this study). Here, we cross-validated the model, therefore, model estimation and evaluation occurred on the same speech stimulus, namely the 12-minute long story. Secondly, we derived acoustic NT for a forward modeling approach, while Decruy et al. (2019) reported acoustic NT with the backward modeling approach. In the backward modeling approach, the stimulus representations are reconstructed from the EEG data rather than the EEG is predicted based on the stimulus representations (the forward modeling approach). We used the forward modeling approach because it gives additional temporal information, i.e., how the brain responds to certain speech characteristics at specific latencies, and spatial distribution, i.e., how these brain responses are spatially distributed across the different EEG electrodes. Such spatial and temporal information is not gained using the backward modeling approach. However, the advantage of the backward model is that it optimally combines information from all EEG electrodes. Therefore, it can regress out non-stimulus-related EEG responses, resulting in more robust reconstruction accuracies. If the effect of age on NT is widespread and subtle, using across-electrode information might be essential to obtain robust results, which would explain why this effect is not observed using the forward modeling approach.

**Figure 4:**
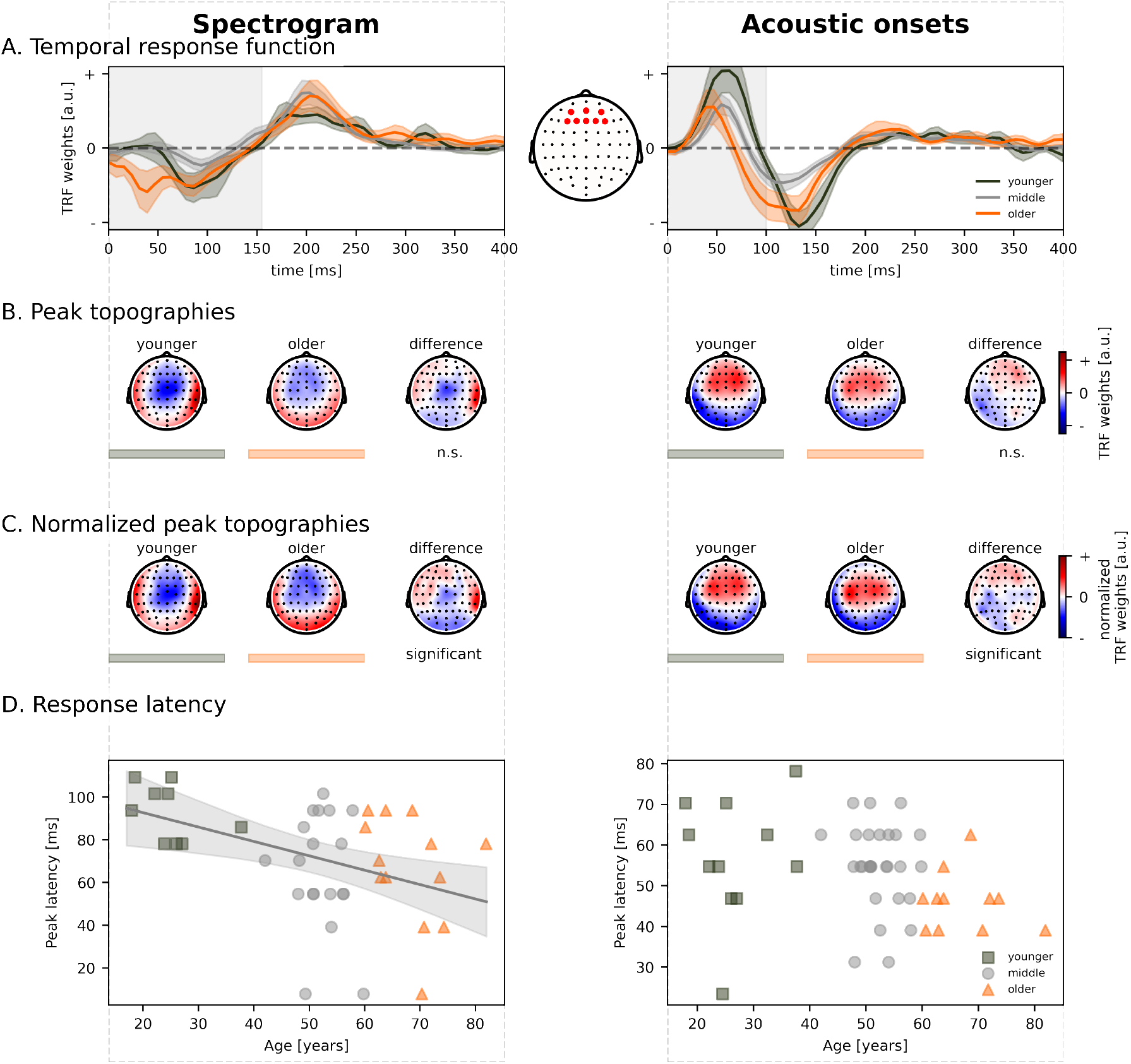
Older adults show shorter latencies for the early acoustic response. The neural responses to acoustic speech representations are visualized as a function of age for both speech representations: spectrogram (left) and acoustic onsets (right). **A.** The average temporal response functions across frontocentral electrodes (indicated on the middle inset) for younger (green), middle-aged (grey), and older (orange) participants. The shaded area indicates the time region used to find the peak topography. **B.** This panel shows the corresponding peak topographies associated with the peak found in the grey vertical panel. **C.** The corresponding normalized peak topographies associated with the peak found in the grey vertical panel. To test whether the normalized topographies significantly differed, the mcCarthy Wood method was applied (for more details see section 2.10). **D.** The decrease in neural response latency as a function of age. The grey lines indicate the model predictions and 95% confidence interval of how the response latency increases with age. The data points of adults above 60 years (older) are annotated with the orange triangles, while those of the adults below 40 years (younger) are annotated with the green squares. Not all subjects showed a prominent peak; therefore, the plot does not show these data points. The insets show the corresponding peak topographies for respectively the younger and older subjects. *n.s. = not significant*

#### 3.2.2 Hemispheric lateralization of neural tracking

Using a linear mixed effects model, a significant age and hemisphere interaction effect was found (interaction effect: p = 0.020). The age effect in the right hemisphere was significant, whereas this was not the case for the left hemisphere (right: p = 0.043; left: p = 0.330), indicating that with increasing age, acoustic information was less strongly represented in the right hemisphere’s speech-evoked neural responses whereas the contribution of the left hemisphere was stable across age (fig. 3.C). In order to identify which spatial pattern may be responsible for this effect, we analyzed the same effect in the 3 ROIs. We found an age and hemisphere interaction effect in the temporal ROI (p = 0.048, FDR-corrected) (fig. 3.D), as well as a significant main effect of age (p = 0.012, FDR-corrected), but no main effect of hemisphere (p = 0.080, FDR-corrected). Post-hoc analyses showed that in the temporal ROI, the age effect in the right hemisphere was significant, whereas this was not the case for the left hemisphere (right: p *<* 0.001; left: p = 0.432). No interaction effects between age and hemisphere were found for the other 2 ROIs.

#### 3.2.3 Neural response latency

Using a linear model, we investigated whether age impacts the latency of the neural responses to acoustic representations. As visualized in fig. 4 and confirmed by the statistical analysis, with increasing age there was a decrease in latency of the early response to the spectrogram representation (p = 0.023, FDR-corrected). We did not observe an effect of age on the later response to both acoustic speech representations nor on the early response to acoustic onsets. The SEM analysis indicated that the observed age effect on the early response latency of the spectrogram was neither mediated through PTA nor cognitive measures (see supplementary material section 6.4.2).

We compared the topographies for the early response to both acoustic representations of the older (*>* 60 years) and younger (*>* 40 years) participants (visualized in fig. 4.B). For the acoustic onsets, the topography of the early response did not significantly differ between older and younger participants. However, for the spectrogram representation, a significant difference was observed (the encircled cluster shows the location which drives this significant difference). After normalization of the topographies, i.e. removing amplitude differences between the two groups, a significant difference between the peak topographies was found between younger and older adults (normalized amplitudes are visualized in fig. 4.C; ANOVA-analysis: age-electrode interaction for the early response to the spectrogram: F(126, 2623) = 1.82, p *<* 0.001; age-electrode interaction for the early response to acoustic onsets: F(126, 3199) = 1.40, p = 0.002).

As a supplementary analysis, we investigated whether the spatial patterns of the TRF amplitudes, i.e. the non-normalized topographies, of the acoustic speech representations averaged across the selected time windows (see table 2) would change across age. The detailed results can be found in the supplementary material (tables S.1, S.2,S.3 and figure S.7). No main effect of age was found on this spatial pattern. This result is aligned with the results of the topography cluster-based permutation test using two discrete age groups.

**Table 3:**
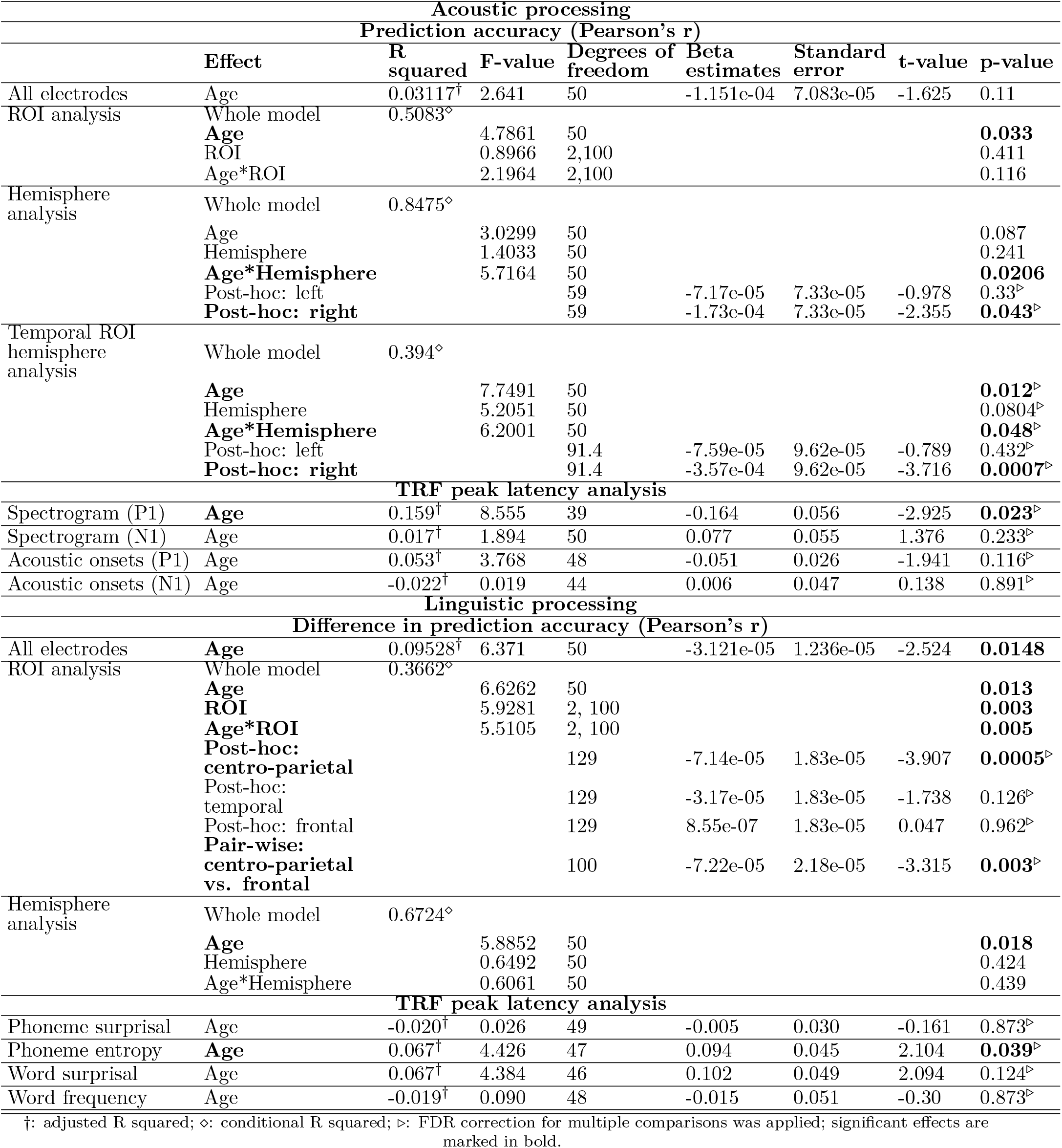
Acoustic and linguistic speech processing results.

Regarding the hemisphere analysis, we found that the TRF weights of the early peak of acoustic onsets show significantly different age trends in the left than in the right hemisphere (see supplementary material section 6.6.1, table S.1 and figure S.7)). This is in line with findings from the prediction accuracy analysis. Analyzing the ROIs, we found no main effect of age, unlike the analysis of the prediction accuracies. Instead, a significant main effect of ROI was observed for the later peaks of the spectrogram and the acoustic onsets TRF. Post-hoc tests revealed that the frontal ROI had the largest contribution to these later peaks. Furthermore, for the early peaks of both speech representations, we found a significant interaction effect between age and ROI. For the early acoustic onsets TRF peak (a positive peak), the frontal ROI showed decreasing TRF weights with advancing age and this age trend was significantly different from the one in the temporal ROI. For the early spectrogram TRF peak (a negative peak), there was a significant difference of the age trend between the centro-parietal ROI (increase with advancing age) and the temporal ROI (decrease with advancing age). Taking into account the opposite polarity of these two early acoustic peaks, together these findings show an increasing contribution of the temporal ROI with advancing age (see supplementary material section 6.6.1).

### 3.3 Effect of age on linguistic speech processing

#### 3.3.1 Prediction accuracy

We repeated the analysis as mentioned earlier for linguistic NT. A measure of linguistic NT is calculated as the model’s prediction accuracy increase when the linguistic representations are included compared to a model without these representations. This is a conservative way to determine the added value of linguistic speech representations: our tests quantify the variability *uniquely* attributable to these representations, not counting variance that is shared between acoustic and lexical segmentation representations. As a first analysis, we determined whether age affects this difference in prediction accuracy averaged across all electrodes and later integration windows (160-300 ms and 300-600 ms). Using a linear model, we observed that linguistic NT decreases with increasing age (p = 0.0148; fig.3.F). Statistics are reported in table 3. A similar trend is observed when testing the individual age groups using cluster-based permutation tests: for younger participants central and temporal channels show significant linguistic tracking (significant model improvement in 4 clusters, p = 0.003) while such improvement is not observed for older adults. The results are shown in supplementary figure S.3.

The linear mixed effects model used to investigate the effect of aging in 3 ROIs showed a significant main effect of age (p = 0.013), ROI (p = 0.003), as well as a significant interaction between age and ROI (p = 0.005). The age effect in the centro-parietal ROI was significant, whereas this was not the case for the other two ROIs (centro-parietal: p = 0.0005; temporal: p = 0.126; frontal: p = 0.962; fig.3.G). This analysis showed that with increasing age, linguistic NT decreases which is most prominent in centro-parietal areas. Pairwise comparisons revealed that the age effect in the centro-parietal ROI is significantly different from the one in the frontal ROI (p = 0.003) while not significantly different from the temporal ROI.

Using SEM analyses, we investigated whether the observed effects are directly due to age or whether the observed age effect in the linear models is spurious due to the correlations between age and cognition, tested via SCWT, RST, or hearing levels, tested via PTA. By investigating the linguistic NT across these two windows for both, across all electrodes and centro-parietal electrodes, this direct effect of age persisted after including predictors for hearing capacity and cognition (more details regarding the SEM analyses in the supplementary material). These analyses suggest that the effect of decreasing linguistic NT with increasing age cannot be attributed to age-related decline in hearing capacity or cognition.

Combining the two windows implies that the linguistic responses at the level of phonemes and words are grouped. By investigating the individual windows, we observed that the linguistic tracking in the window from 160 to 300 ms, targeting the N250 response to linguistic representations at the phoneme level, was not affected by age. However, in the later window, i.e., 300 to 600 ms, the effect of age was mediated through SCWT. Moreover, investigating this effect in the different ROIs suggested that frontal areas drive this effect (see supplementary material section 6.5).

#### 3.3.2 Hemispheric lateralization of neural tracking

Same as for the acoustic model we were interested in hemispheric lateralization changes with age. However, no significant age and hemisphere interactions were found, neither for the whole hemispheres, nor for any of the ROIs. The main effect of hemisphere was also not significant, only the main effect of age was significant (p = 0.018).

#### 3.3.3 Neural response latency to linguistic speech representations

We observed a difference in TRF between younger and older adults for the linguistic speech representations. The peaks around 250 ms and 400 ms for respectively the linguistic representations at the phoneme and word level tend to be more prominent for younger adults, while these peaks show higher variability in latency and amplitude among older adults (fig. S.4 and fig. S.5). However, statistical testing showed that only the latency of the N250 response to phoneme entropy showed a significant age effect after correcting for multiple comparisons (FDR correction with n=4), i.e., longer latencies were observed for older participants (p = 0.039, see table 3). No such age effect was observed for the other three linguistic representations. Interestingly, the SEM analysis showed that this increase in latency of the N250 peak to phoneme entropy showed an indirect effect of age, mediated through hearing capacity. Adults with a worse hearing showed a longer latency of the N250 response to phoneme entropy.

Surprisingly, the average TRFs of older adults do not show a significant peak around 400 ms (see supplementary figures S.4 and S.5). We attribute this to two reasons: firstly, the amplitude of the older participants in this time window is smaller, and the variability in spatial and temporal pattern of the TRF is higher across older participants. Although the amplitude is small, we could still determine a peak latency as we identified a peak based on its template topography. Within a specific time region (table 1), we searched for a topography resembling the template topography at each peak of the TRF, identified by the RMS value of the TRF as specified in section *Determination of peak latency*.

As the SEM analysis suggested that frontal areas might be impacted differently by SCWT, we investigated the TRFs in the frontal channel selection. For these TRFs, we observed more prominent negative activity for older adults. Therefore, we investigated for each age category whether the TRF was significantly negative to determine whether or not a robust negative peak is seen (annotated by the lines above the TRFs in fig. 5). For phoneme surprisal, phoneme entropy and word surprisal, we only observed significant negative clusters for middle-aged and older adults. For word frequency, a significant negative cluster is observed for younger and older participants (see table 4). Additionally, we compared whether the TRF is significantly more negative for older adults compared to younger adults. This was the case for phoneme surprisal (cluster from 55 to 70 ms, p = 0.0428, corrected for 4 comparisons) and word surprisal (cluster from 141 to 156 ms, p = 0.048, corrected for 4 comparisons).

We compared the topographic responses of the older (*>* 60 years) and younger (*<* 40 years) participants (visualized as insets in fig. S.4, panels B and C). We did not observe a significant difference in the peak topographies (using the permutation test) nor a difference in the normalized peak topographies (using the McCarthy Wood method).

**Figure 5:**
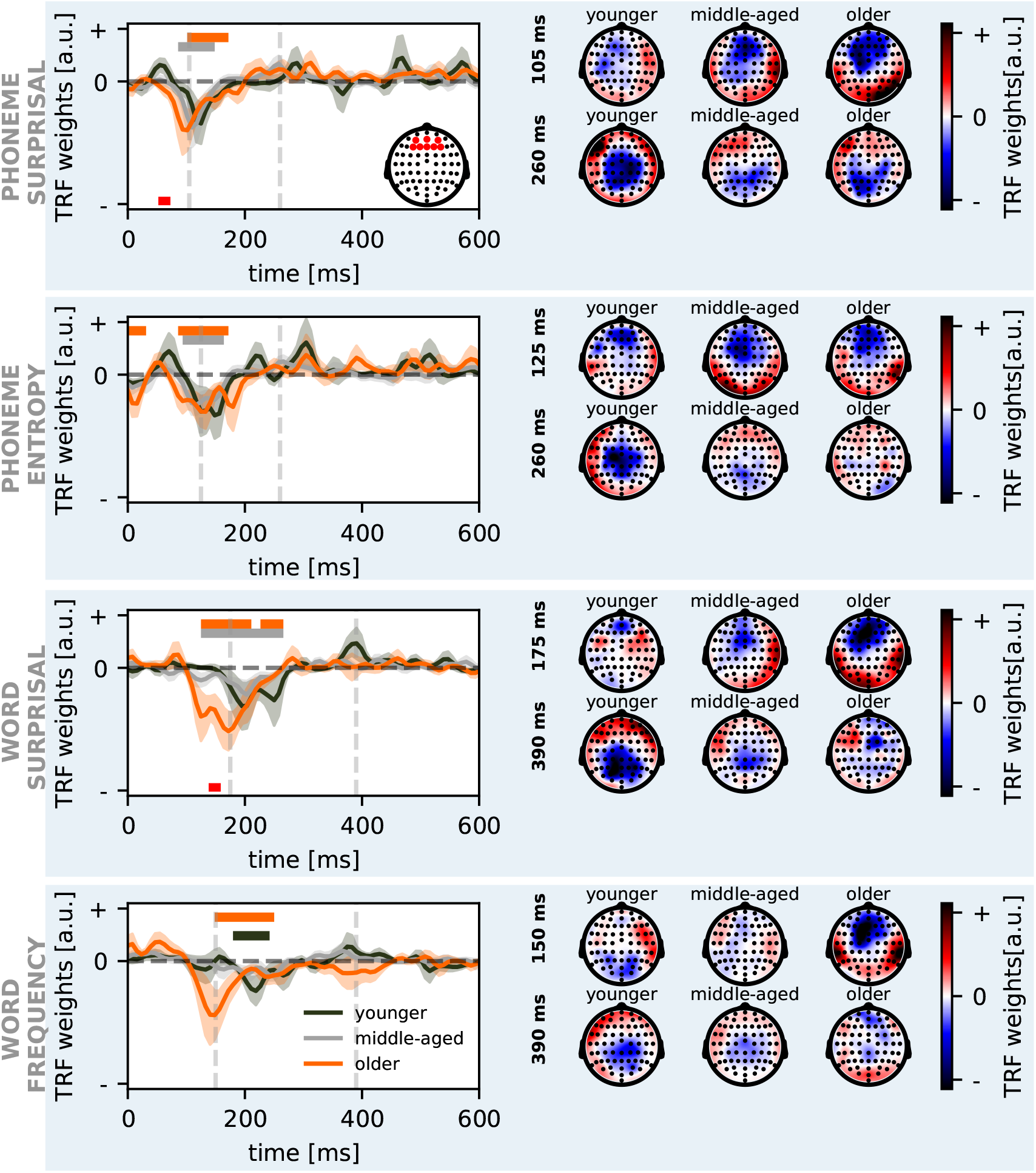
Older adults show an enhanced response to linguistic representations in fronto-central regions. The neural responses to linguistic representations are visualized in function of age for the three linguistic speech representations: phoneme surprisal, (top), phoneme entropy, word surprisal and word frequency (bottom). The left plot shows the average temporal response functions across frontocentral electrodes (indicated on the inset of the top plot) for younger (green), middle (grey), and older-aged (orange) participants. The shaded area indicates the standard error of the average TRF. The lines above the TRFs indicate where the TRF is significantly negative for each age category (annotated in the corresponding color). The red bold lines underneath the TRFs indicate where the TRFs are significantly different between younger and older adults. The right-sided plots show the corresponding peak topographies associated with the peaks at the time of the dashed grey line in the left-sided plots.

**Table 4:** Observed significant negative clusters in the linguistic TRFs in frontocentral regions for middle-aged and older adults.

| feature | clusters observed<br>in middle-age adults | clusters observed<br>in older adults |
| --- | --- | --- |
| phoneme surprisal | 86 to 148 ms ($p = 0.011$) | 102 to 172 ms ($p < 0.001$) |
| phoneme entropy | 94 to 164 ms ($p = 0.003$) | 0 to 31 ms ($p = 0.033$)<br>86 to 172 ms ($p < 0.001$) |
| word surprisal | 125 to 266 ms ($p < 0.001$) | 125 to 211 ms ($p < 0.001$)<br>227 to 266 ms ($p = 0.029$) |

Furthermore, we also analyzed the spatial patterns of the TRF amplitudes of the linguistic speech representations, i.e., phoneme surprisal, phoneme entropy, word surprisal and word frequency averaged across the selected time windows (see table 2). The detailed results can be found in the supplementary material (tables S.2, S.3 and figure S.7). The hemisphere analysis revealed no significant effects, which is in line with results from the unique contribution of linguistic speech representations. The ROI analysis revealed a main effect of ROI for each of the speech representations, but only in the later TRF peaks (between 250 and 450ms). At these negative peaks, the TRF weights in the centro-parietal ROI were significantly lower than in the frontal and temporal ROI. This is partly in line with the results of the analysis of the unique contribution of linguistic speech representations in that the centro-parietal ROI seems to be contributing the most to this peak. However, the interaction effect between age and ROI was not significant for any of the speech representations after applying correction for multiple comparisons.

## 4 Discussion

In order to investigate whether and how linguistic and acoustic speech processing change from early to late adulthood, we analyzed how the brain tracks respective speech representations when listening to natural speech. We modelled the unique contribution of linguistic speech representations while controlling for acoustic processing. Spatial information was investigated by analyzing a priori defined ROIs, by analyzing hemispheric lateralization and by investigating TRF topographies across age. Temporal patterns of speech integration were investigated by analyzing TRF latencies. SEM was applied to account for the influence of hearing ability and cognition on NT of natural speech.

Age affected both acoustic and linguistic speech processing. With increasing age, neural tracking generally decreased. Spatially, older adults relied less on the right hemisphere for acoustic processing, which was most prominent in temporal regions. Although no age-related change in hemispheric preference was observed for linguistic speech processing, linguistic NT decreased with increasing age, which was driven by centro-parietal regions. Exploring temporal patterns of the neural response, we found that older adults displayed earlier peaks to acoustic cues while they displayed later peaks to linguistic aspects of speech. In this normal-hearing population sample, we could not attribute these age-related differences in speech processing (both acoustic and linguistic) to an age-related decline in hearing or cognition. However, there might be some influence of cognition on the neural response latency of word-level linguistic speech processing.

### 4.1 Age effects on tracking of acoustic features

#### 4.1.1 Acoustic neural tracking decreases with increasing age

This study shows that acoustic NT is not affected by age when considering all electrodes. However, when averaging over frontal, temporal and centro-parietal electrodes, acoustic NT decreases with advancing age (fig. 3.B). Hitherto, speech envelope tracking studies investigating aging trends have reported results opposing the current ones, namely increased acoustic NT with increasing age (Presacco et al., 2016; Brodbeck et al., 2018b; Decruy et al., 2019; Karunathilake et al., 2020). This is, however, likely due to methodological differences, e.g., a different modelling approach (backward versus forward modelling), different acoustic speech representations (envelope versus spectrogram and acoustic onsets), different paradigms or a different composition of the studied sample (age categories versus age as a continuous variable).

When this effect of age on acoustic NT is further investigated, we observed that hearing capacity affects acoustic tracking in the opposite direction. Namely, individuals with decreasing hearing capacity, associated with higher PTA, show higher acoustic tracking. This finding is consistent with literature investigating the impact of hearing loss on acoustic neural tracking (Decruy et al., 2020; Fuglsang et al., 2020). Although it is difficult to disentangle the impact of age or hearing loss on neural responses, our results suggest that both processes alter neural responses differently. Moreover, this age effect was not mediated through cognitive measures.

By investigating hemispheric lateralization of acoustic NT, we found that with increasing age, participants rely less on the right hemisphere for acoustic speech processing. This effect was most prominent in temporal regions. The finding that older adults show a left-hemispheric preference agrees with the findings by Brodbeck et al. (2018b). Brodbeck et al. (2018b) suggested that higher acoustic NT with increasing age might not necessarily imply enhanced encoding of acoustic representations but might indicate increased neural activity in the left temporal lobe. Likewise, Farahani et al. (2020) found enhanced activity in older adults in the left hemisphere for processing simple, non-speech, acoustic stimuli with modulations at the syllable rate. This poses the question whether this left-hemispheric preference can also be linked to higher-order auditory processing. Although the current study cannot give a definite answer, our results suggest that this is unrelated to working memory and inhibition.

#### 4.1.2 Older adults show earlier responses to acoustic representations

The temporal pattern analysis unveiled shorter TRF latencies with increasing age for the early peak of the spectrogram (fig. 4.A & C). Based on previous literature using simple sounds as clicks, tones, or syllables, we expected to find an age difference in the later acoustic response (N1 and P2, around 100-200ms) rather than the early acoustic response (before 100ms). In general, we hypothesized longer latencies with advancing age, but the current results demonstrate the opposite. One possible explanation might be the task demand: listening to speech is an automatic process that humans are highly trained at, whereas processing artificially created clicks, tones or syllables is less familiar and might thus influence the neural response differently. Moreover, neural speech processing is modulated by attention effects. Older adults may be more involved with the task than younger adults, resulting in shorter neural response latencies to acoustic speech processing. Using word pairs, Roque et al. (2019) also observed a decrease in early acoustic response latency in older adults.

By investigating the normalized topographies associated to these early acoustic responses, we observed a significant difference between the youngest and oldest adults (fig. 4.B). This finding supports that the neural activity underlying this early acoustic response is different between younger and older adults. This is not surprising as with increasing age, the brain significantly changes in structure and function (Salthouse, 2011; Voineskos et al., 2012; Reuter-Lorenz and Cappell, 2008; Tubi et al., 2020). As we did not perform source localization, it is difficult to say how this neural activation is different. Using source localization, Brodbeck et al. (2018b) reported that older adults recruited additional regions at a short latency, around 30 ms, relative to the slow amplitude modulations of the speech envelope.

### 4.2 Age effects on linguistic processing

#### 4.2.1 Linguistic neural tracking decreases with increasing age

In this study, we show that linguistic NT decreases with increasing age. Specifically, older adults showed a lower neural tracking magnitude than younger adults in response to four linguistic speech representations, i.e., phoneme surprisal, phoneme entropy, word surprisal and word frequency. These findings deviate from previously reported results, i.e., Broderick et al. (2021) and Mesik et al. (2021). Broderick et al. (2021) did not find a significant difference between age groups in NT of word surprisal, while Mesik et al. (2021) reported a higher neural tracking for older compared to younger adults. However, it is not straightforward to compare our results to those previously published results due to substantial methodological differences. Firstly, both studies relied on a reduced set of stimulus features, therefore not considering acoustic representations. Controlling for acoustics is important as the speech acoustics can explain apparent responses to higher-level speech representations (Daube et al., 2019), leading to biased results (Gillis et al., 2021). Secondly, the speech was presented in a different format. While in the current study and the study by Broderick et al. (2021), a single-speaker paradigm was used, Mesik et al. (2021) investigated the neural responses to an attended and ignored speaker. As a dual-talker paradigm is more effortful, this change in paradigm might influence the neural responses. Thirdly, the determination of word surprisal differs among the 3 studies. The current study and Broderick et al. (2021) relied on a simple 5-gram model which determines the word surprisal given a context window of the 4 previous words. Mesik et al. (2021) relied on a more complex language model, i.e. GTP-2, which relies on broader context windows. Lastly, also the age-distribution differs across these studies. Broderick et al. (2021) and Mesik et al. (2021) investigated two age groups, while in the current study, we investigated age as a continuous variable. Moreover, the current study, as well as Broderick et al. (2021), relied on more participants of older age compared to Mesik et al. (2021).

Using the SEM analysis, we observed that this effect of age on linguistic NT, averaged across two time windows, was not mediated by hearing capacity nor cognitive measures. However, these time windows characterize different type of processing, 160 to 300 ms captures the N250 response at the level of phonemes and 300 to 600 ms captures the N400 response, a word level linguistic response. Therefore, we investigated the individual time windows. No effect of age was observed on the window capturing linguistic processing at the phoneme level. However, we observed that this age effect only persists for the time window from 300 to 600 ms, where presumably word-level processing occurs. Interestingly, we observed that linguistic tracking, averaged across all channels in this later window, showed an indirect effect of age mediated through the SCWT score, i.e., older adults with a better cognitive performance show higher linguistic tracking. Breaking this effect down across the different ROIs, we observed that this effect is driven by the frontal areas. This finding agrees with known age-related neurophysiological changes in the brain (Peelle et al., 2010), i.e, older adults have a reduced ability to recruit regions in the left ventral inferior frontal gyrus. Although, our findings agree in the sense that we observed a reduced activity in frontal areas, we did not observe a left-hemispheric lateralization. In sum, our results show a complex interplay between age, linguistic NT and cognition, suggesting that cognition plays an important role in the prominence of the age-related decline of linguistic NT.

#### 4.2.2 Early frontal neural response in older adults

Older adults, but not younger adults showed a negative TRF peak of phoneme and word surprisal over frontal electrodes between 100 and 200 ms. We assume that this peak is related to another neural response than the one typically associated with word surprisal around 400ms. With the current dataset, we cannot attribute a specific mechanism to this effect, however one explanation might be age-related neurophysiological changes in the brain.

#### 4.2.3 Older adults display later neural responses to phoneme entropy

The neural response latency to the phoneme entropy peak between 250 and 350ms increased with advancing age. This means that older adults need more processing time to capture the information represented by phoneme entropy. At a domain-general cognitive level, earlier research has reported decreased processing speed in older adults (Salthouse, 1996; Salami et al., 2012; Salthouse, 2011; Bennett and Madden, 2014). Surprisingly, the SEM analysis suggested that the age effect is mediated through hearing capacity but not cognitive measures. Such a mediation effect might indicate that distorted bottom-up processes affect these linguistic processes. We can only speculate what the increase in neural response latency at phoneme level processing in older adults means, given that phoneme-level linguistic representations have not yet been investigated in older adults.

### 4.3 Caveats

Similar to most studies investigating neural tracking, this study too has the caveat that we rely on the assumption that the speech is fully understood by the participants. We believe this assumption is valid as the speech was presented in silence and the behavioural measures that were collected confirm this. However, behaviourally measuring understanding of continuous natural speech is challenging.

Another caveat of the current study is that results regarding the hemispheric lateralization might be affected by the right-lateralized stimulus presentation. We are not aware how these results generalize to a similar dataset where the stimulus is presented bilaterally.

Furthermore, since we found decreasing acoustic NT across ROIs with advancing age, the degraded acoustic input may hamper the interaction between bottom-up acoustic and top-down linguistic speech processing aspects. This way, the lower-fidelity auditory input may propagate onto higher-level processes.

Another limitation is that here, we only had outcome measures of 2 cognitive subdomains available, i.e., inhibition and working memory. It would be good in the future to also take into account other cognitive subdomains when assessing the role of cognition in language processing across the adult lifespan.

The last caveats of the current study relate to the approach to determine linguistic NT. A first limitation is that we relied on the feature set determined by Gillis et al. (2021). Although they investigated which linguistic representations contributed unique information on top of acoustic representations, they do not guarantee that this feature set is a sufficient set, i.e., covers all aspects of linguistic language processing, nor that this feature set is optimal to use in older adults. A second caveat is the limited amount of data. All participants listened to the same 12-minute-long story. Twelve minutes is not much data to train all the used speech representations. Previous literature investigating linguistic representations relied on 30 minutes or more data. Therefore, certain effects might not be observed in the current dataset due to the limited amount of data. It would be interesting to investigate from what amount of data on the neural tracking of different speech representations would stabilize in normal-hearing, cognitively unaffected older adults in the future using a similar approach as e.g., Di Liberto and Lalor (2017). The dataset used in the current study does not allow to investigate this as 12 minutes was not sufficient to obtain a significant response in older adults. Notice that the first two caveats might explain why no significant linguistic tracking was observed in older adults. Given that no linguistic tracking was observed, one should interpret the linguistic TRFs cautiously as these results’ uncertainty is high. Third, we controlled for a large part of the acoustic variance in the linguistic representations by using 4 acoustic and segmentation-related speech representations. However, we did not control for all acoustic aspects that might be captured by linguistic features, e.g. pitch or F0. Thus, linguistic NT might still contain some lower-level processing activity. Also, the current approach, i.e., investigating the added value of linguistic representations, assumes that the shared variance between acoustic and linguistics is attributed to acoustics. This approach is, therefore, somewhat restrictive as it looks at the unique information contributed by the linguistic representations on top of the acoustic information. However, the current approach does not allow for a more depth investigation of the shared variance between acoustics and linguistics. Moreover, we did not account for the variance in the acoustic NT that is explained by linguistic information. Lastly, by relying on linear methods as we did here, the brain’s non-linearity is not accounted for. Therefore, it may be interesting to explore this research questions in the future via non-linear methods, such as mutual information (Zan et al., 2020).

## 5 Conclusion

This study illustrates that both acoustic and linguistic speech processing mechanisms change across the adult lifespan, which is underpinned by spatiotemporal changes. For the first time, the unique contribution of NT of linguistic speech processing was investigated in older adults. We demonstrated that linguistic speech processing declines with advancing age. Moreover, as age increased, the neural response latency to certain aspects of linguistic speech processing increased. These age-related changes were mainly apparent over centro-parietal electrodes. Acoustic NT decreased as well with increasing age, but in contrast to linguistic processing older subjects showed shorter latencies for the early acoustic response to speech. Different neural resources were involved in acoustic speech processing in older than in younger adults, e.g., less involvement of the right hemisphere with advancing age.

Furthermore, we observed that linguistic speech processing in older adults might be affected by a complex interplay between age-related structural changes and functional changes which might be modulated by cognition at the word-level. In future studies analyzing neural speech tracking across the adult lifespan, it would be interesting to explore this complex interplay by taking into account additional behavioral correlates of cognition and by using source localization and functional connectivity methods in experiments using natural, continuous speech.

Our results indicate that linguistic neural tracking decreases across the lifespan. This study shows the need to investigate further how linguistic mechanisms change during aging to find a robust measure of speech comprehension in older adults. Such a measure of linguistic processing would significantly impact the diagnosis and speech therapy of people with language and communication disorders, such as aphasia, which predominantly affect older adults.

## Data Availability Statement

After completion of the review process the preprocessed data, i.e. the neural tracking values, neural response latencies, etc., will be made available. The code for the template matching is available via the GitHub repository (https://github.com/exporl/TemplateMatching).

## Funding Sources

The presented study received funding from the European Research Council (ERC) under the European Union’s Horizon 2020 research and innovation programme (Tom Francart; grant agreement No. 637424). Research of Marlies Gillis (PhD grant: SB 1SA0620N) was funded by the Research Foundation Flanders (FWO). Research of Jill Kries was supported by the Luxembourg National Research Fund (FNR) (AFR-PhD project reference 13513810). Furthermore, this study was financially supported by the FWO, grant No. G0D8520N.

## Disclosures

No conflicts of interest, financial or otherwise, are declared by the authors.

## Acknowledgements

The authors would like to thank Dr. Toivo Glatz for methodological advice, Prof. Dr. Astrid Van Wieringen for advice on the cognitive part of the data and Dr. Klara Schevenels for providing valuable comments on the introduction and discussion. We would also like to thank Dr. Lien Decruy and Dr. Jonas Vanthornhout for collecting the dataset used in this study. They were assisted in data collection by Elien Van den Borre, Melissa Schoubben, Dr. Annelies Devesse, and Dr. Sam Denys. A thank you also goes to all participants.

## Author contributions

TF, JK, MG and MV conceived and designed research; JK and MG analyzed data; JK, MG, TF and MV interpreted results of experiments; JK and MG prepared figures; JK and MG drafted manuscript; MV, TF, MG and JK edited and revised manuscript; MV, TF, MG and JK approved final version of manuscript.

## 6 Supplementary Material

### 6.1 Pure tone audiometry

**Figure S.1:**
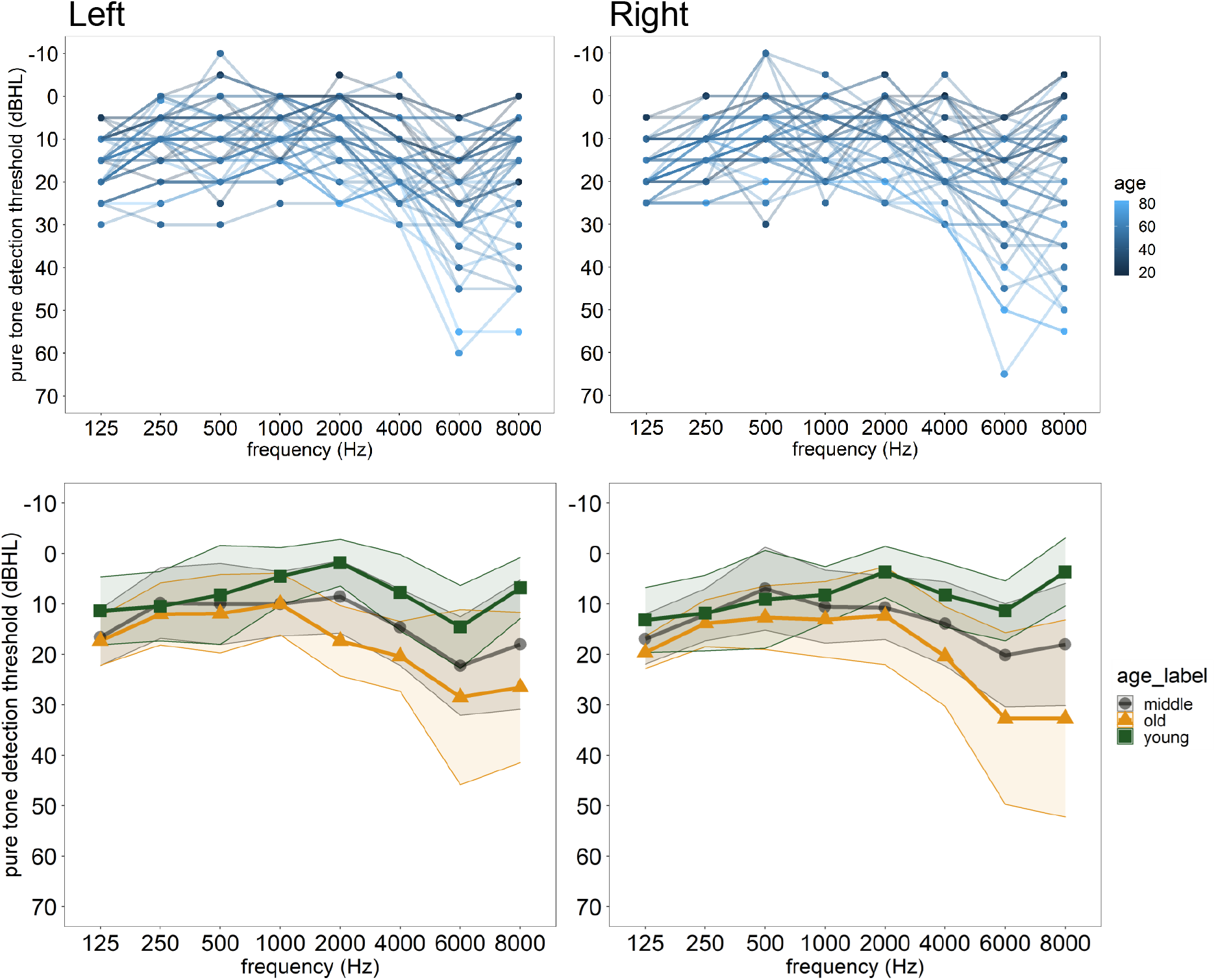
Visualization of the pure tone audiogram thresholds. The top panels show the raw data for the left and the right ear. The bottom panels show the mean and standard deviation of the younger, middle-aged and older group for the left and the right ear.

### 6.2 Neural tracking of lexical segmentation: phoneme and word onsets

We visualized the TRFs to phoneme and word onsets in fig. S.2 for younger, middle-aged, and older participants. Next to this analysis, we compared how including the lexical segmentation features affects the TRFs to the acoustic features (not visualized). An almost perfect overlap was found between the TRFs for the spectrogram and acoustic edges obtained with models with and without the lexical segmentation features.

**Figure S.2:**
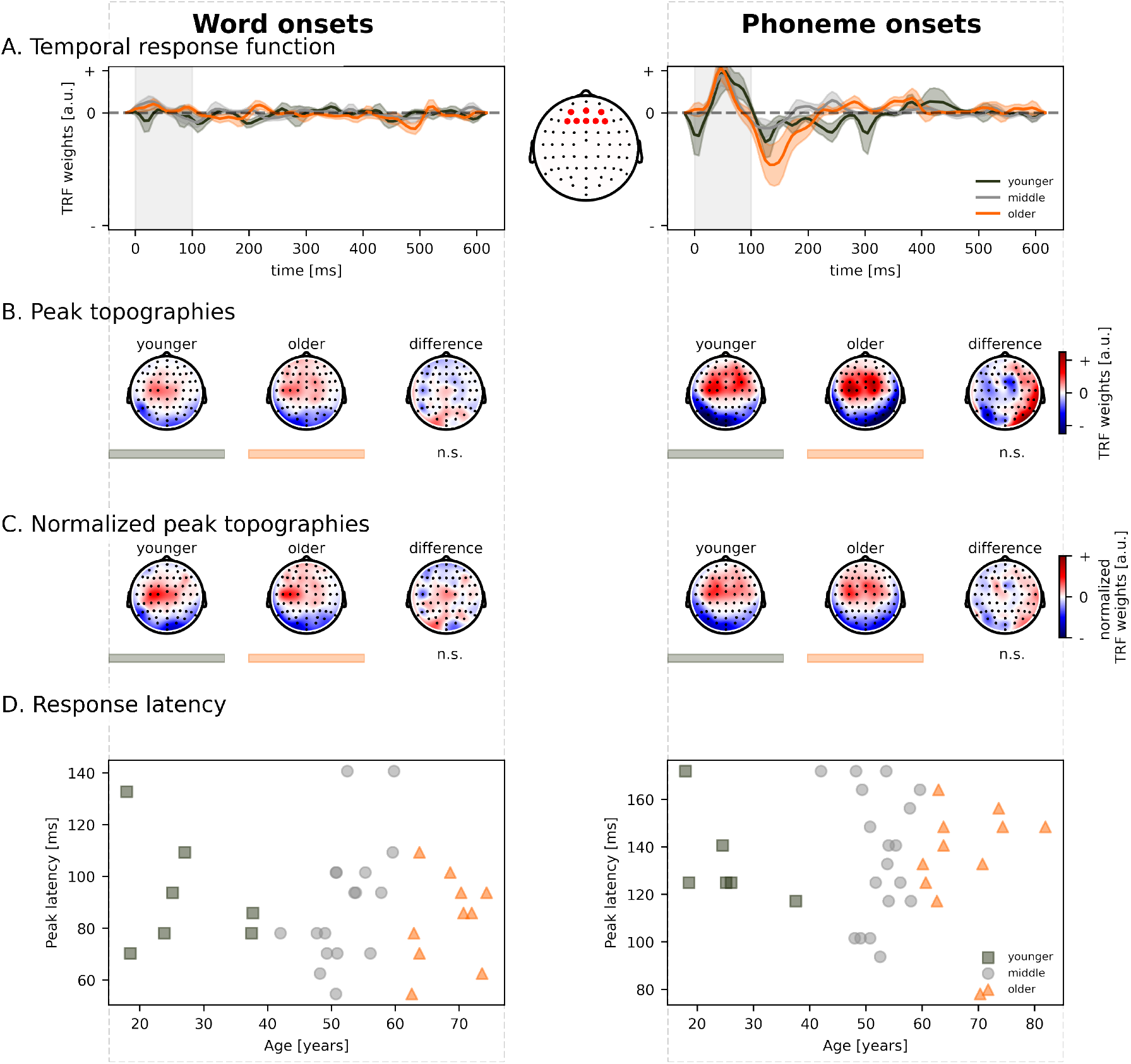
No differences across age in the neural processing of lexical segmentation. The neural responses to lexical segmentations are visualized as a function of age for two speech representations: word onsets (left) and phoneme onsets (right). **A.** The average temporal response functions across frontal electrodes (indicated on the middle inset) for younger (green), middle-aged (grey), and older (orange) participants. **B.** The corresponding peak topographies associated with the peak found in the grey vertical panel. **C.** The corresponding normalized peak topographies associated with the peak found in the grey vertical panel. **D.** The neural response latency across age. The data points of adults above 60 years (older) are annotated with the orange triangles, while those of the adults below 40 years (younger) are annotated with the green squares. Not all subjects showed a prominent peak; therefore, the plot does not show these data points. *n.s. = not significant*

### 6.3 Neural tracking of linguistic features

**Figure S.3:**
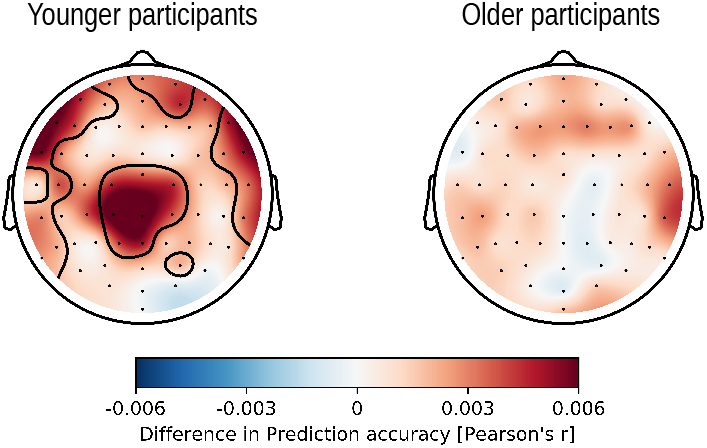
Linguistic tracking in younger (left) and older (right) adults. Results of the cluster-based permutation test on the added value of linguistic representations. The clusters which drive the topography to be significantly higher than 0 are encircled. For younger adults, different clusters are observed while no clusters were observed for older adults.

### 6.4 Structural Equation Modelling – Acoustics

#### 6.4.1 Neural Tracking

Using these standardized estimates, we applied SEM to assess the effect of PTA and age on the prediction accuracy averaged across the three ROIs and across the two time windows: 30-160 ms and 160-300 ms. We observed a marginal indirect effect of age through the hearing capacity (represented by PTA) on the acoustic NT (model specifications: RMSEA = 0, CFI = 1, TLI = 1.035; saturated model; indirect effect; *β* = 0.156, SE = 0.077, z-value = 2.016, p = 0.044). The PTA is affected by age (*β* = 0.478, SE = 0.122, z-value = 3.927, p *<* 0.001), and PTA affects the acoustic NT (*β* = 0.325, SE = 0.138, z-value = 2.349, p = 0.019). We observed both a direct effect of age on acoustic NT: as age increases, acoustic NT decreases (direct effect; *β* = -0.502, SE = 0.166, z-value = -3.020, p = 0.003). A schematic overview of the SEM results is visualized in the left panel of fig. S.6.

Therefore, even though the degree of hearing loss is taken into account, a direct effect of age on acoustic neural tracking is still observed. Interestingly, the indirect effect of age has an opposite sign than the direct age effect. As age increases, the degree of hearing loss increases reflected by higher PTA values which increases the neural tracking. The latter assumption converges with studies investigating the impact of hearing loss: hearing-impaired listeners show an enhanced neural tracking of speech compared to normal-hearing listeners (Decruy et al., 2020; Fuglsang et al., 2020).

**Figure S.4:**
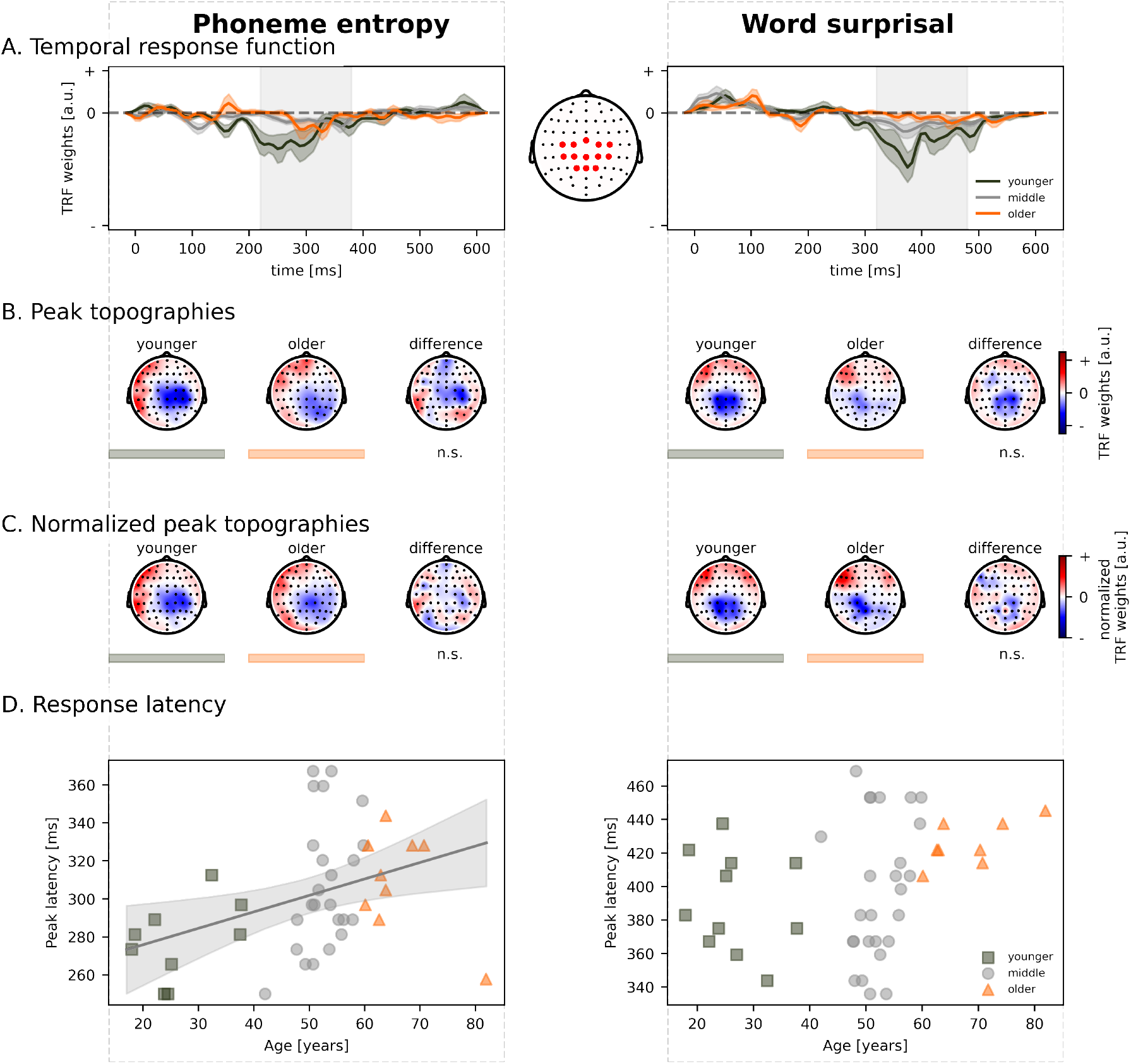
Older adults show longer latencies for linguistic speech processing. The neural responses to linguistic representations are visualized as a function of age for two linguistic speech representations: phoneme entropy (left) and word surprisal (right). **A.** The average temporal response functions across central electrodes (indicated on the middle inset) for younger (green), middle-aged (grey), and older (orange) participants. **B.** The corresponding peak topographies associated with the peak found in the grey vertical panel. **C.** The corresponding normalized peak topographies associated with the peak found in the grey vertical panel. **D.** The increase in neural response latency across age. The grey lines indicate the model predictions and 95% confidence interval of how the response latency increases with age. The neural response latency across age. The data points of adults above 60 years (older) are annotated with the orange triangles, while those of the adults below 40 years (younger) are annotated with the green squares. Not all subjects showed a prominent peak; therefore, the plot does not show these data points. *n.s. = not significant*

**Figure S.5:**
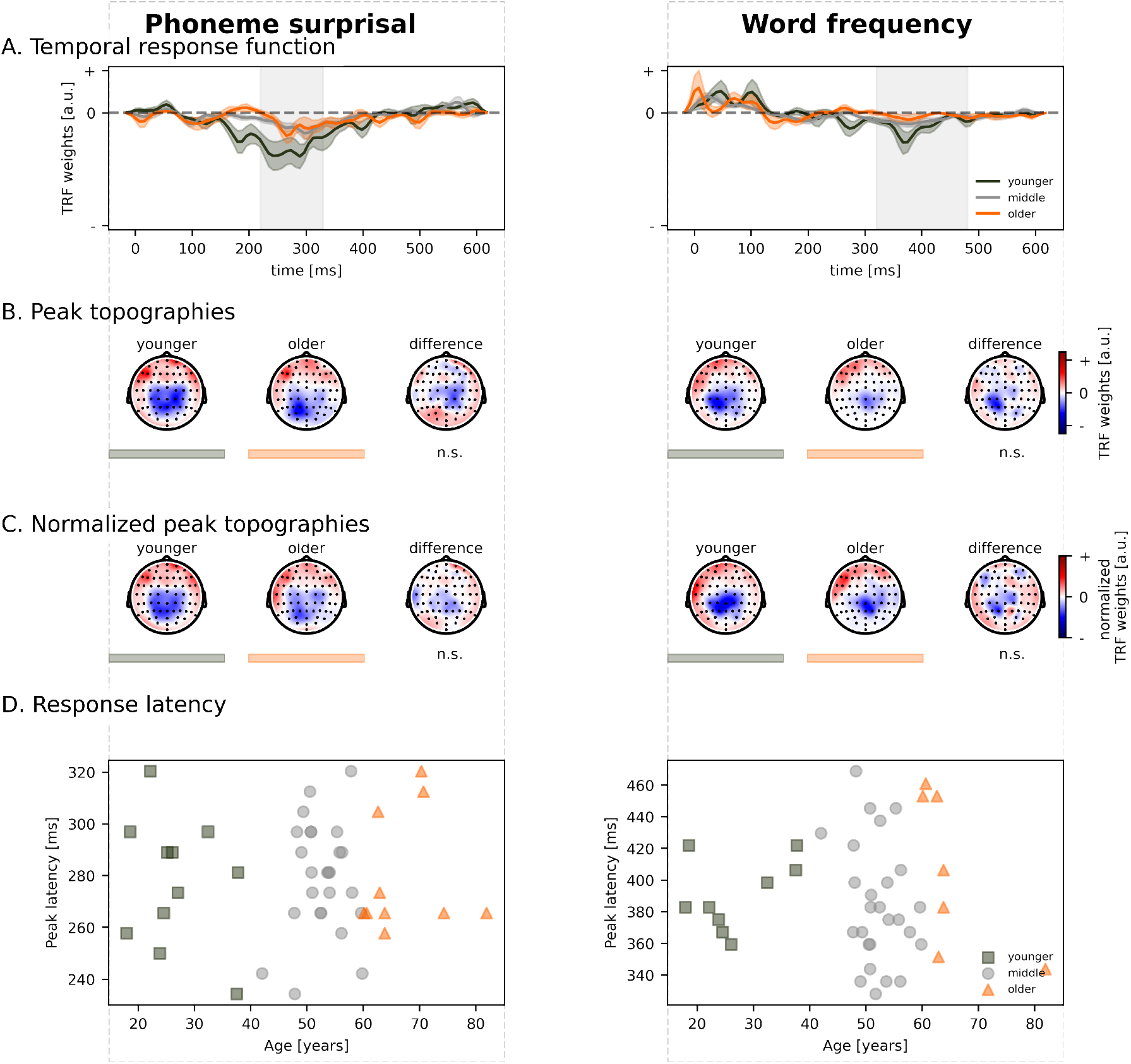
No differences across age in the neural processing of phoneme surprisal and word frequency. The neural responses to lexical segmentations are visualized as a function of age for two linguistic speech representations: phoneme surprisal (left) and word frequency (right). **A.** The average temporal response functions across central electrodes (indicated on the middle inset) for younger (green), middle-aged (grey), and older (orange) participants. **B.** The corresponding peak topographies associated with the peak found in the grey vertical panel. **C.** The corresponding normalized peak topographies associated with the peak found in the grey vertical panel. **D.** The neural response latency across age. The data points of adults above 60 years (older) are annotated with the orange triangles, while those of the adults below 40 years (younger) are annotated with the green squares. Not all subjects showed a prominent peak; therefore, the plot does not show these data points. *n.s. = not significant*

#### 6.4.2 Neural Response Latency

We investigated the SEM for each peak latency for each speech representation separately. As age is highly correlated with the degree of hearing loss, SEM allows us to model the effect of age directly onto the response latency as well as the effect of age mediated through the PTA values. The effect of age was not mediated through PTA for any peak latency. More details regarding these analyses are given in the following paragraphs (schematic overview of the SEM results are visualized in fig. S.6). As only for the early response to the spectrogram, a significant effect of age was observed, we further investigated this age effect on the neural response latency of this peak.

For the response latency of the first peak of the spectrogram (30 to 125 ms), we observed a direct effect of age onto the response latency using a linear model. This direct effect of age did not reach significance in the SEM analysis (model specifications: RMSEA = 0, CFI = 1, TLI = 1; saturated model; direct age effect; *β* = -0.122, SE = 0.063, z-value = -1.947, p = 0.051). Although age significantly impacts the PTA (*β* = 0.495, SE = 0.128, z-value = 3.872, p *<* 0.001), there is no effect of PTA on the latency of this first peak in the spectrogram (*β* = -0.085, SE = 0.066, z-value = -1.296, p =0.195). Therefore, we do not observe a significant indirect effect of age (indirect age effect; *β* = -0.042, SE = 0.034, z-value = -1.947, p = 0.219). Although the indirect or direct age effects do not reach significance, the total effect of age did reach significance (*β* = -0.164, SE = 0.055, z-value = -2.999, p = 0.003). This suggests that the SEM analysis cannot disentangle the direct and indirect effect on the neural response latency of the spectrogram.

### 6.5 Structural Equation Modelling – Linguistic

#### 6.5.1 Neural Tracking

To investigate whether a decrease in hearing capacity or cognition explained the age effect observed across all electrodes and two later windows, we applied SEM. However, across all electrodes, we did not observe a mediation effect of age through SCWT, RST, or PTA (model specifications: RMSEA = 0, CFI = 1, TLI = 1.029). Age impacts linguistic NT directly (direct age effect: *β* = -0.365, SE = 0.158, z-value = -2.316, p = 0.021). Age also significantly impacts the SCWT (*β* = 0.493, SE = 0.121, z-value = 4.086, p *<* 0.001), RST (*β* = -0.443, SE = 0.124, z-value = -3.565, p *<* 0.001), and PTA (*β* = 0.478, SE = 0.122, z-value = 3.921, p *<* 0.001). SCWT significantly mediated linguistic NT, while RST and PTA did not (effect of SCWT on linguistic NT: *β* = -0.379, SE = 0.133, z-value =-2.857, p = 0.004; effect of RST on linguistic NT: *β* = -0.071, SE = 0.129, z-value = -0.554, p = 0.579; effect of PTA on linguistic NT: *β* = 0.212, SE = 0.132, z-value = 1.613, p = 0.107). This results in a total effect of age on linguistic NT is *β* = -0.419 (SE = 0.127, z-value = -3.294, p = 0.001). This indicates that age explains % of the observed total effects of age (i.e. including the indirect age effects mediated through the SCWT, RST and PTA).

**Figure S.6:**
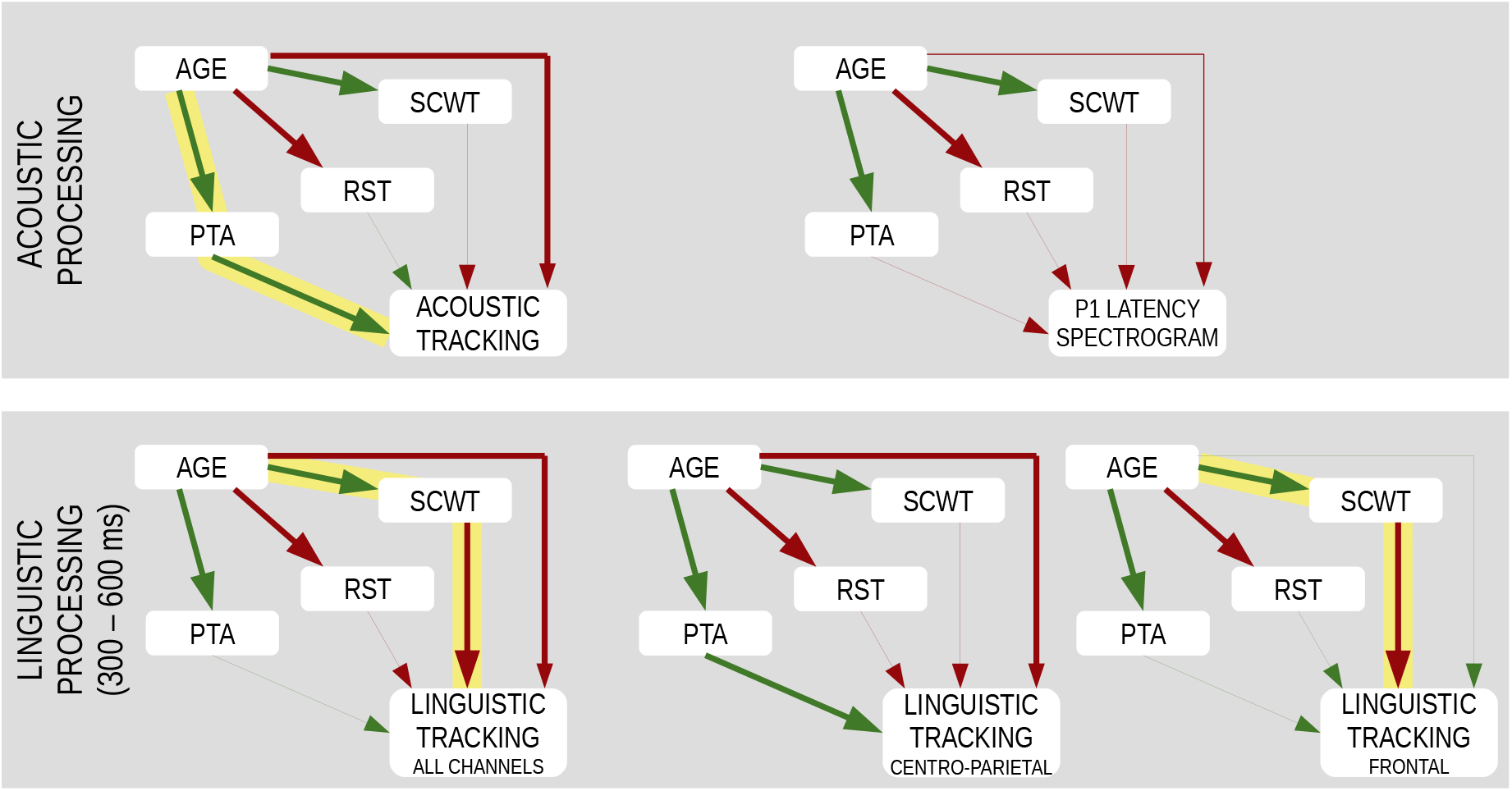
Results obtained with SEM for the acoustic model for neural tracking (top, left panel; for the channels included in the ROIs as indicated on fig. 1) and latency of the early neural responses to the spectrogram (top, right panel). The bottom panel shows the results of the SEM analysis using the time window from 300 to 600 ms across all channels (left), centro-parietal (middle) and frontal channels (right). Positive effects are indicated in a green arrow, while negative effects are shown in red. Significant effect are annotated with a bold arrow. The indirect effects of age are annotated by the yellow marker.

When the linguistic NT values are averaged across centro-parietal electrodes and these two later time windows, the effect of age is not mediated through SCWT, RST, or PTA (model specifications: RMSEA = 0, CFI = 1, TLI = 1.032). Here, the direct age effect of age is *β* = -0.522 (SE = 0.164, z-value = -3.176, p = 0.001), while the total effect of age, i.e. direct and indirect age effects, is *β* = -0.466 (SE = 0.123, z-value = -3.788, p *<* 0.001).

In the later time window (300 to 600 ms), we observed significant impact of age on linguistic NT when averaged across all electrodes and the centro-parietal electrodes channel selection, but not when averaged across the frontal nor the temporal electrodes electrode selection (using a linear modeling approach). To investigate these effects in more detail, we applied SEM. For simplicity purposes, we fitted three models: (1) a SEM based on the linguistic NT values across all electrodes, (2) a SEM across centro-parietal electrodes, and (3) a SEM across frontal and temporal electrodes. These results are visualized in the left panel of fig. S.6.

However, in the late time window (300-600ms), averaged across all electrodes, we observed a direct age effect and an indirect effect of age through the SCWT scores (model specifications: RMSEA = 0, CFI = 1, TLI = 1.029). We observed a direct effect of age on linguistic NT (*β* = -0.365, SE = 0.158, z-value = -2.316, p = 0.021). No indirect effect of PTA and RST were observed. However, we did observe a significant indirect effect of age through the SCWT score(*β* = -0.187, SE = 0.080, z-value = -2.342, p = 0.019): the SCWT score increases with increasing age (*β* = 0.493, SE = 0.121, z-value = 4.086, p *<* 0.001), and a higher SCWT score decreases linguistic NT (*β* = -0.379, SE = 0.133, z-value = -2.857, p = 0.004). The total indirect effect of age on linguistic NT is *β* = -0.054 (SE = 0.120, z-value = -0.452, p =0.652; including the effect of SCWT, RST, and PTA). Altogether, this suggests that the age explains 87% of the observed total effects of age.

For the centro-parietal electrode selection, the effect of age on linguistic NT was not mediated by PTA, RST or SCWT (model specifications: RMSEA = 0, CFI = 1, TLI = 1.026). We observed a direct age effect (*β* = -0.639, SE = 0.148, z-value = -4.308, p *<* 0.001). However, no indirect effect of age through PTA, RST or SCWT reached significance. Although the effect of PTA on linguistic NT reached significance (*β* = 0.257, SE = 0.124, z-value = 2.074, p = 0.038), the mediating effect of age through PTA was not significant (*β* = 0.123, SE = 0.067, z-value = 1.833, p = 0.067). The total indirect of effect of age is *β* = 0.067 (SE = 0.107, z-value = 0.624, p = 0.532). The direct age affect therefore explains most of the observed total age effect.

The indirect age effect mediated through SCWT scores was only observed when the prediction accuracies were averaged across all electrodes. As this effect was absent in the centro-parietal electrode selection, we also analysed the frontal and temporal electrode selection. Indeed, in the frontal electrode selection, although we did not observe a significant direct age effect, we observed a significant indirect effect of age mediated through the SCWT scores (model specifications: RMSEA = 0, CFI = 1, TLI = 1.039; indirect effect of SCWT: *β* = -0.179, SE = 0.086, z-value = -2.084, p = 0.037). The direct age effect becomes non-significant, which align with the results of the above analysis (*β* = 0.105, SE = 0.178, z-value = 0.593, p = 0.553). With increasing age, the SCWT scores increase (*β* = 0.493, SE = 0.121 z-value = 4.086, p *<* 0.001), which leads to a decrease in frontal linguistic NT (*β* = -0.362, SE = 0.150, z-value =-2.423, p = 0.015).

#### 6.5.2 Neural Response Latency

Using linear modelling, we observed that increasing age was associated with longer response latencies of the N250 response to phoneme entropy. To gain more insight in this effect, we investigate whether this age effect was mediated through cognitive measures or hearing capacity. The SEM analysis showed a direct effect of age on the N250 response latency of phoneme entropy (model specifications: RMSEA = 0.05, CFI = 0.99, TLI = 0.97; direct age effect; *β* = 0.129, SE = 0.050, z-value = 2.597, p = 0.009). Moreover, we observed a significant indirect age effect mediated through hearing capacity (*β* = -0.051, SE = 0.025, z-value = -2.026, p = 0.043). Interestingly, the indirect effect of age mediated through hearing capacity is opposite compared to the direct age effect.

### 6.6 Hemisphere and ROI analysis of the TRF amplitude in selected time windows

The spatial analysis (i.e., hemisphere and ROI analysis) that was conducted on the magnitude of neural tracking was also performed for the TRF weights in selected time windows. These results are reported in tables S.1, S.2 and S.3. The results are visualized in figure S.7.

**Figure S.7:**
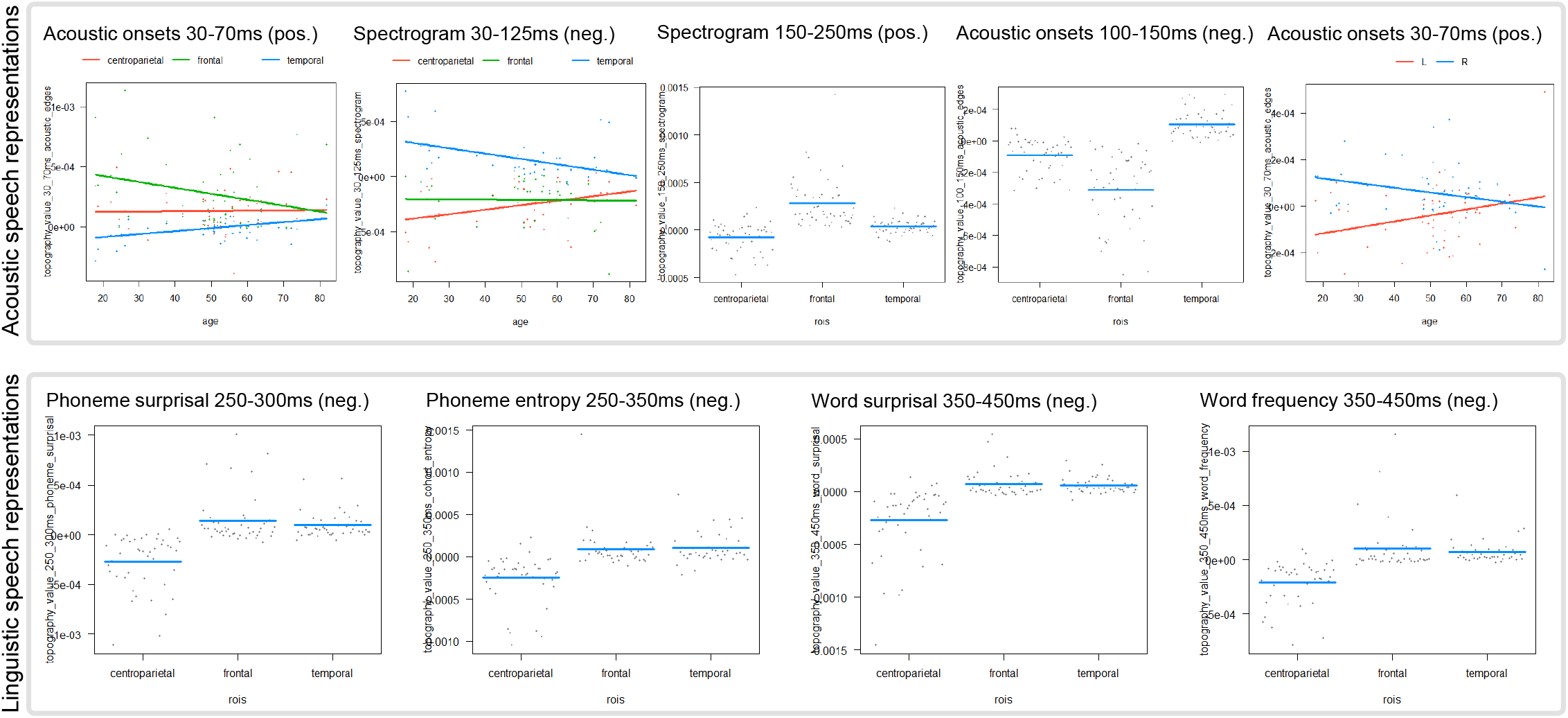
Visualization of the results of the TRF topography analysis. See following pages for more details.

#### 6.6.1 Summary of findings: Acoustic speech representations

##### Hemisphere analysis

- No main effect of age in neither of the two peaks of the spectrogram nor the two peaks of acoustic onsets.
- Significant main effect of hemisphere for the early peak (30-70ms) of acoustic onsets, which is a positive peak.
- Significant interaction effect of age and hemisphere also only for the early peak (30-70ms) of acoustic onsets (positive peak). The TRF weights in the left hemisphere showed a significantly different age trend than the right hemisphere (left: increasing with advancing age; right: decreasing with advancing age).

##### ROI analysis

- No main effect of age across ROIs in neither of the two peaks of the spectrogram nor the two peaks of acoustic onsets.
- Significant main effect of ROI in all four peaks. In the late peaks of both the spectrogram (positive peak betwwen 150 and 250ms) and acoustic onsets (negative peak between 100 and 150ms), the frontal ROI had the largest contribution to those TRF peaks.
- The early peaks of both speech representations (spectrogram: 30-125ms = negative peak; acoustic onsets: 30-70ms = positive peak) showed significant interaction effects between age and ROI, in addition to the significant main effect of ROI. The post-hoc tests for the interaction effect of the negative spectrogram peak showed that the age effect was significant for the centro-parietal (increasing with advancing age) and the temporal (decreasing with advancing age) ROIs and that the TRF weights in the centro-parietal ROI had a different age trend than in the temporal ROI. The post-hoc tests for the interaction effect of the positive acoustic onsets peak showed that the TRF weights in the frontal ROI were significant and that this age trend (i.e., decreasing with advancing age) is significantly different from the age trend in the temporal ROI.

**Table S.1:** Spatial analysis of the TRFs of acoustic speech representations.

| Analysis | Effect | F-value | Degrees of freedom | Beta estimates | Standard error | t-value | p-value |
| --- | --- | --- | --- | --- | --- | --- | --- |
| **Spectrogram: 30-125ms** |  |  |  |  |  |  |  |
| Hemisphere | Age | 0.2774 | 1, 78 |  |  |  | 0.8962 |
|  | Hemisphere | 4.0401 | 1, 78 |  |  |  | 0.0957 |
|  | Age*Hemisphere | 0.7343 | 1, 78 |  |  |  | 0.5254 |
| ROI | Age | 0.0936 | 1, 117 |  |  |  | 0.8713 |
|  | **ROI** | 19.4318 | 2, 117 |  |  |  | **<0.0001** |
|  | **Age\*ROI** | 5.7019 | 2, 117 |  |  |  | **0.0173** |
| Post-hoc | **centro-parietal** |  | 117 | 4.09e-06 | 1.88e-06 | 2.180 | **0.0469** |
|  | **temporal** |  | 117 | -4.87e-06 | 1.88e-06 | -2.595 | **0.032** |
|  | **centro-parietal - temporal** |  | 78 | 8.95e-06 | 2.65e-06 | 3.376 | **0.0034** |
| **Spectrogram: 150-250ms** |  |  |  |  |  |  |  |
| Hemisphere | Age | 0.0171 | 1, 100 |  |  |  | 0.8962 |
|  | Hemisphere | 0.9966 | 1, 100 |  |  |  | 0.4273 |
|  | Age*Hemisphere | 1.1559 | 1, 100 |  |  |  | 0.5254 |
| ROI | Age | 0.1592 | 1, 150 |  |  |  | 0.8713 |
|  | **ROI** | 6.1586 | 2, 150 |  |  |  | **0.0026** |
|  | **Age\*ROI** | 1.0723 | 2, 150 |  |  |  | 0.4597 |
| Post-hoc | **centro-parietal** |  | 150 | -7.86e-05 | 2.41e-05 | -3.268 | **0.002** |
|  | **frontal** |  | 150 | 2.81e-04 | 2.41e-05 | 11.668 | **<0.0001** |
|  | **centro-parietal - frontal** |  | 100 | -0.000359 | 3.4e-05 | -10.562 | **<0.0001** |
|  | **centro-parietal - temporal** |  | 100 | -0.000119 | 3.4e-05 | -3.494 | **0.0007** |
|  | **frontal - temporal** |  | 100 | 0.000240 | 3.4e-05 | 7.068 | **<0.0001** |
| **Acoustic onsets: 30-70ms** |  |  |  |  |  |  |  |
| Hemisphere | Age | 0.1197 | 1, 96 |  |  |  | 0.8962 |
|  | **Hemisphere** | 15.1579 | 1, 96 |  |  |  | **0.0007** |
|  | **Age\*Hemisphere** | 8.042 | 1, 96 |  |  |  | **0.0222** |
| Post-hoc | **left - right** |  | 48 | 4.57e-06 | 1.61e-06 | 2.836 | **0.0067** |
| ROI | Age | 0.4369 | 1, 144 |  |  |  | 0.8713 |
|  | **ROI** | 11.2338 | 2, 144 |  |  |  | **<0.0001** |
|  | **Age\*ROI** | 4.0371 | 2, 144 |  |  |  | **0.0393** |
| Post-hoc | **frontal** |  | 144 | -4.93e-06 | 1.9e-06 | -2.597 | **0.0311** |
|  | **frontal - temporal** |  | 96 | -7.44e-06 | 2.68e-06 | -2.771 | **0.0201** |
| **Acoustic onsets: 100-150ms** |  |  |  |  |  |  |  |
| Hemisphere | Age | 0.5781 | 1, 88 |  |  |  | 0.8962 |
|  | Hemisphere | 0.2242 | 1, 88 |  |  |  | 0.637 |
|  | Age*Hemisphere | 0.2987 | 1, 88 |  |  |  | 0.586 |
| ROI | Age | 0.0263 | 1, 132 |  |  |  | 0.8713 |
|  | **ROI** | 10.4947 | 2, 132 |  |  |  | **<0.0001** |
|  | **Age\*ROI** | 0.378 | 2, 132 |  |  |  | 0.6859 |
| Post-hoc | **centro-parietal** |  | 132 | -8.89e-05 | 2.26e-05 | -3.939 | **<0.0001** |
|  | **temporal** |  | 132 | 1.08e-04 | 2.26e-05 | 4.792 | **<0.0001** |
|  | **frontal** |  | 132 | -3.12e-04 | 2.26e-05 | -13.805 | **<0.0001** |
|  | **centro-parietal - frontal** |  | 88 | 0.000223 | 3.19e-05 | 6.977 | **<0.0001** |
|  | **centro-parietal - temporal** |  | 88 | -0.000197 | 3.19e-05 | -6.174 | **<0.0001** |
|  | **frontal - temporal** |  | 88 | -0.000420 | 3.19e-05 | -13.150 | **<0.0001** |

All p-values were corrected for multiple comparisons (n=4, except for post-hoc tests n=2 (hemisphere) or n=3 (ROIs)) using the FDR. Significant effects are marked in bold. Only significant post-hoc tests are reported.

#### 6.6.2 Summary of findings: (Pre-)lexical speech representations

##### Hemisphere analysis

- Significant main effect of hemisphere and interaction effect of age and hemisphere at the early peak of phoneme onsets, i.e., a positive peak between 30 and 70ms.
- The post-hoc analyses showed that the age trend in both hemispheres was significant (left: increased with advancing age; right: decreased with advancing age) and that the age trends between the right and the left hemisphere differed from each other.
- Significant main effect of hemisphere for the late peak of phoneme onsets (100-150ms), i.e. a negative peak. The left hemisphere TRF weights were significantly more negative than the right hemisphere.

##### ROI analysis

- No significant effects.

**Table S.2:** Spatial analysis of the TRFs of (pre-)lexical speech representations.

|  | Effect | F-value | Num<br>DF | Den<br>DF | Estimate | SE | t-value | P-value |
| --- | --- | --- | --- | --- | --- | --- | --- | --- |
| phoneme_onsets_30_70ms |  |  |  |  |  |  |  |  |
| Hemisphere | age | 0.054522312 | 1 | 94 |  |  |  | 0.815880813 |
|  | **hemisphere** | 9.683714664 | 1 | 94 |  |  |  | **0.009850091** |
|  | **age:hemisphere** | 11.71833714 | 1 | 94 |  |  |  | **0.003673206** |
| Post-hoc | **left** |  |  | 94 | 5.70E-06 | 2.53E-06 | 2.255462 | **0.02642** |
|  | **right** |  |  | 94 | -6.54E-06 | 2.53E-06 | -2.585681 | **0.0225** |
|  | **left - right** |  |  | 47 | 1.22E-05 | 3.58E-06 | 3.423206 | **0.0012** |
| ROI | age | 0.135584453 | 1 | 141 |  |  |  | 0.791897463 |
|  | rois | 0.40856248 | 2 | 141 |  |  |  | 0.665389177 |
|  | age:rois | 1.196576113 | 2 | 141 |  |  |  | 0.701825311 |
| phoneme_onsets_100_150ms |  |  |  |  |  |  |  |  |
| Hemisphere | age | 0.487006812 | 1 | 66 |  |  |  | 0.650289216 |
|  | **hemisphere** | 5.647997615 | 1 | 66 |  |  |  | **0.040768754** |
|  | **age:hemisphere** | 2.920060372 | 1 | 66 |  |  |  | 0.18436719 |
| Post-hoc | **left - right** |  |  | 33 | -0.000131 | 4.97E-05 | -2.636 | **0.0127** |
| ROI | age | 0.289413447 | 1 | 99 |  |  |  | 0.791897463 |
|  | rois | 2.335527382 | 2 | 99 |  |  |  | 0.408277104 |
|  | age:rois | 0.307422669 | 2 | 99 |  |  |  | 0.736779997 |
| word_onsets_30_70ms |  |  |  |  |  |  |  |  |
| Hemisphere | age | 1.174444998 | 1 | 78 |  |  |  | 0.563654689 |
|  | hemisphere | 0.070963994 | 1 | 78 |  |  |  | 0.790641293 |
|  | **age:hemisphere** | 0.011508785 | 1 | 78 |  |  |  | 0.914843098 |
| ROI | age | 0.906061721 | 1 | 117 |  |  |  | 0.791897463 |
|  | rois | 1.049509359 | 2 | 117 |  |  |  | 0.665389177 |
|  | age:rois | 0.306264849 | 2 | 117 |  |  |  | 0.736779997 |
| word_onsets_100_120ms |  |  |  |  |  |  |  |  |
| Hemisphere | age | 1.298848484 | 1 | 66 |  |  |  | 0.563654689 |
|  | hemisphere | 0.32853861 | 1 | 66 |  |  |  | 0.757958312 |
|  | **age:hemisphere** | 0.12650488 | 1 | 66 |  |  |  | 0.914843098 |
| ROI | age | 0.069992668 | 1 | 99 |  |  |  | 0.791897463 |
|  | rois | 0.504961058 | 2 | 99 |  |  |  | 0.665389177 |
|  | age:rois | 1.058373892 | 2 | 99 |  |  |  | 0.701825311 |

All p-values were corrected for multiple comparisons (n=4, except for post-hoc tests n=2) using the FDR. Significant effects are marked in bold. Only significant post-hoc results are reported. SE=standard error; DF = degrees of freedom.

#### 6.6.3 Summary of findings: Linguistic speech representations

##### Hemisphere analysis

- For the four linguistic speech representations, i.e., phoneme surprisal, phoneme entropy, word surprisal and word frequency, there were no significant main or interaction effects of age and hemisphere.

##### ROI analysis

- For all 4 linguistic speech representations, a main effect of ROI was found, but only in the later TRF peaks (between 250 and 450 ms depending on the speech representation). These late peaks correspond to negative peaks.
- The post-hoc tests revealed that for each of the 4 speech representations, the centro-parietal ROI displayed the largest contribution to these late negative peaks.

**Table S.3:** Spatial analysis of the TRFs of linguistic speech representations.

|  | Effect | F-value | Num<br>DF | Den<br>DF | Estimate | SE | t-value | P-value |
| --- | --- | --- | --- | --- | --- | --- | --- | --- |
| **phoneme_surprisal_80_150ms** |  |  |  |  |  |  |  |  |
| Hemisphere | age | 0.2544 | 1 | 80 |  |  |  | 0.7551 |
|  | hemisphere | 5.7277 | 1 | 80 |  |  |  | 0.1281 |
|  | age:hemisphere | 3.3226 | 1 | 80 |  |  |  | 0.5765 |
| ROI | age | 0.0918 | 1 | 120 |  |  |  | 0.8713 |
|  | rois | 2.1268 | 2 | 120 |  |  |  | 0.1978 |
|  | age:rois | 0.0589 | 2 | 120 |  |  |  | 0.9427 |
| **phoneme_surprisal_250_300ms** |  |  |  |  |  |  |  |  |
| Hemisphere | age | 0.0979 | 1 | 86 |  |  |  | 0.7551 |
|  | hemisphere | 0.0621 | 1 | 86 |  |  |  | 0.8036 |
|  | age:hemisphere | 0.4903 | 1 | 86 |  |  |  | 0.7770 |
| ROI | age | 1.5133 | 1 | 129 |  |  |  | 0.8073 |
|  | **rois** | 8.9443 | 2 | 129 |  |  |  | **0.0006** |
|  | age:rois | 1.2317 | 2 | 129 |  |  |  | 0.5852 |
| Post-hoc | **centro-parietal** |  |  | 129 | -0.00028 | 3.4068E-05 | -8.39542 | **2.148E-13** |
|  | **frontal** |  |  | 129 | 0.00013 | 3.4068E-05 | 4.03675 | **0.0001** |
|  | **temporal** |  |  | 129 | 9.5816E-05 | 3.4068E-05 | 2.81245 | **0.0056** |
|  | **centro-parietal - frontal** |  |  | 86 | -0.00042 | 4.8180E-05 | -8.79087 | **3.935E-13** |
|  | **centro-parietal - temporal** |  |  | 86 | -0.00038 | 4.8180E-05 | -7.92516 | **1.124E-11** |
| **phoneme_entropy_100_175ms** |  |  |  |  |  |  |  |  |
| Hemisphere | age | 0.1758 | 1 | 72 |  |  |  | 0.7551 |
|  | hemisphere | 0.0972 | 1 | 72 |  |  |  | 0.8036 |
|  | age:hemisphere | 0.0402 | 1 | 72 |  |  |  | 0.9618 |
| ROI | age | 0.4322 | 1 | 108 |  |  |  | 0.8196 |
|  | rois | 1.3185 | 2 | 108 |  |  |  | 0.3623 |
|  | age:rois | 0.8769 | 2 | 108 |  |  |  | 0.5852 |
| **phoneme_entropy_250_350ms** |  |  |  |  |  |  |  |  |
| Hemisphere | age | 0.1302 | 1 | 80 |  |  |  | 0.7551 |
|  | hemisphere | 4.7621 | 1 | 80 |  |  |  | 0.1281 |
|  | age:hemisphere | 0.5689 | 1 | 80 |  |  |  | 0.7770 |
| ROI | age | 2.7923 | 1 | 120 |  |  |  | 0.7785 |
|  | **rois** | 10.633 | 2 | 120 |  |  |  | **0.0002** |
|  | age:rois | 3.4424 | 2 | 120 |  |  |  | 0.1407 |
| Post-hoc | **centro-parietal** |  |  | 120 | -0.00027 | 3.7313E-05 | -7.27019 | **1.196E-10** |
|  | **frontal** |  |  | 120 | 8.1421E-05 | 3.7313E-05 | 2.18206 | **0.0310** |
|  | **temporal** |  |  | 120 | 0.00010 | 3.7313E-05 | 2.89460 | **0.0067** |
|  | **centro-parietal - frontal** |  |  | 80 | -0.00035 | 5.2769E-05 | -6.68376 | **4.282E-09** |
|  | **centro-parietal - temporal** |  |  | 80 | -0.00037 | 5.2769E-05 | -7.18760 | **9.261E-10** |
| **word_surprisal_125_225ms** |  |  |  |  |  |  |  |  |
| Hemisphere | age | 0.1188 | 1 | 72 |  |  |  | 0.7551 |
|  | hemisphere | 2.0647 | 1 | 72 |  |  |  | 0.4135 |
|  | age:hemisphere | 0.0003 | 1 | 72 |  |  |  | 0.9860 |
| ROI | age | 0.1803 | 1 | 35.9992 |  |  |  | 0.8713 |
|  | rois | 0.6656 | 2 | 71.9992 |  |  |  | 0.5170 |
|  | age:rois | 0.5362 | 2 | 71.9992 |  |  |  | 0.6711 |
| **word_surprisal_350_450ms** |  |  |  |  |  |  |  |  |
| Hemisphere | age | 0.2773 | 1 | 86 |  |  |  | 0.7551 |
|  | hemisphere | 0.7792 | 1 | 86 |  |  |  | 0.7596 |
|  | age:hemisphere | 0.2532 | 1 | 86 |  |  |  | 0.8214 |
| ROI | age | 0.7019 | 1 | 129 |  |  |  | 0.8073 |
|  | **rois** | 13.775 | 2 | 129 |  |  |  | **3.025E-05** |
|  | age:rois | 3.5742 | 2 | 129 |  |  |  | 0.140711926 |
| Post-hoc | **centro-parietal** |  |  | 129 | -0.00028 | 3.1417E-05 | -9.00858 | **7.140E-15** |
|  | **frontal** |  |  | 129 | 7.7791E-05 | 3.1417E-05 | 2.47610 | **0.0218** |
|  | **temporal** |  |  | 129 | 6.3469E-05 | 3.1417E-05 | 2.02024 | **0.0454** |
|  | **centro-parietal - frontal** |  |  | 86 | -0.00036 | 4.4430E-05 | -8.12089 | **9.036E-12** |
|  | **centro-parietal - temporal** |  |  | 86 | -0.00034 | 4.4430E-05 | -7.79855 | **2.023E-11** |
| **word_frequency_125_225ms** |  |  |  |  |  |  |  |  |
| Hemisphere | age | 0.4747 | 1 | 80 |  |  |  | 0.7551 |
|  | hemisphere | 0.1856 | 1 | 80 |  |  |  | 0.8036 |
|  | age:hemisphere | 0.6457 | 1 | 80 |  |  |  | 0.7770 |
| ROI | age | 0.0003 | 1 | 120 |  |  |  | 0.9850 |
|  | rois | 0.7532 | 2 | 120 |  |  |  | 0.5170 |
|  | age:rois | 0.8290 | 2 | 120 |  |  |  | 0.58527 |
| **word_frequency_350_450ms** |  |  |  |  |  |  |  |  |
| Hemisphere | age | 1.5698 | 1 | 80 |  |  |  | 0.7551 |
|  | hemisphere | 0.3484 | 1 | 80 |  |  |  | 0.8036 |
|  | age:hemisphere | 1.3290 | 1 | 80 |  |  |  | 0.7770 |
| ROI | age | 0.8051 | 1 | 120 |  |  |  | 0.8073 |
|  | **rois** | 8.6668 | 2 | 120 |  |  |  | **0.0006** |
|  | age:rois | 1.9087 | 2 | 120 |  |  |  | 0.4073 |
| Post-hoc | **centro-parietal** |  |  | 120 | -0.00022 | 2.9487E-05 | -7.48074 | **4.034E-11** |
|  | **frontal** |  |  | 120 | 9.9883E-05 | 2.9487E-05 | 3.38730 | **0.0014** |
|  | **temporal** |  |  | 120 | 7.3389E-05 | 2.9487E-05 | 2.488817 | **0.0141** |
|  | **centro-parietal - frontal** |  |  | 80 | -0.00032 | 4.1701E-05 | -7.68487 | **1.002E-10** |
|  | **centro-parietal - temporal** |  |  | 80 | -0.00029 | 4.1701E-05 | -7.04954 | **8.546E-10** |

All p-values were corrected for multiple comparisons (n=8, except for post-hoc tests n=3) using the FDR. Significant effects are marked in bold. Only significant post-hoc results are reported. SE=standard error; DF = degrees of freedom.

## Notes

https://github.com/MarliesG/TemplateMatching

### Competing Interest Statement

The authors have declared no competing interest.

### Summary of Updates

This version of the manuscript has been revised to update the interpretation of results in the discussion, to clarify some methodological details and to improve the consistency of terminology.

